# Reconstructing the Origins of a Neuropeptide Signaling System Using the Accelerated Evolution of Biodiverse Cone Snail Venoms

**DOI:** 10.1101/2021.11.05.463867

**Authors:** Thomas Lund Koch, Iris Bea L. Ramiro, Paula Flórez-Salcedo, Ebbe Engholm, Knud Jørgen Jensen, Kevin Chase, Baldomero M. Olivera, Walden Emil Bjørn-Yoshimoto, Helena Safavi-Hemami

**Affiliations:** Department of Biomedical Sciences, University of Copenhagen, Copenhagen-N, 2200, Denmark; Department of Neurobiology and Anatomy, University of Utah, Salt Lake City, UT 84112, USA; Department of Chemistry, University of Copenhagen, Frederiksberg, 1871, Denmark; School of Biological Sciences, University of Utah, Salt Lake City UT 84112, USA; Department of Biochemistry, University of Utah, Salt Lake City UT 84112, USA

## Abstract

Somatostatin and its related peptides (SSRPs) form an important family of hormones with diverse physiological roles. The ubiquitous presence of SSRPs in vertebrates and several invertebrate deuterostomes suggests an ancient origin of the SSRP signaling system. However, the existence of SSRP genes outside of deuterostomes has not been established and the evolutionary history of this signaling system remains poorly understood. Our recent discovery of SSRP-like toxins (consomatins) in venomous marine cone snails (*Conus)* suggested the presence of a homologous signaling system in mollusks and potentially other protostomes. Here we identify the molluscan SSRP-like signaling gene that gave rise to the consomatin family. Following recruitment into venom, consomatin genes experience strong positive selection and repeated gene duplications resulting in the formation of a hyper-diverse family of venom peptides. Intriguingly, the largest number of consomatins was found in worm-hunting species (> 400 sequences), indicating a homologous system in annelids, another large protostome phylum. Comprehensive sequence mining enabled the identification of orthologous SSRP-like sequences (and their corresponding orphan receptor) in annelids and several other protostome phyla. These results establish the existence of SSRP-like sequences in many major branches of bilaterians, including xenacoelomorphs, a phylum believed to have emerged before the divergence of protostomes and deuterostomes, ~ 600 My ago. Finally, having a large set of predator-prey SSRP sequences available, we show that while the cone snail’s signaling SSRP-like genes are under purifying selection, in striking contrast, the consomatin genes experience rapid directional selection to target receptors in a changing mix of prey.

## Introduction

Small, secreted peptides form a large and diverse group of signaling molecules that are essential for fundamental biological processes. Many of these signaling peptides originated early in animal evolution with descendants found in both protostomes and deuterostomes, the two major groups of bilaterians (Jékely 2013; Mirabeau and Joly 2013). Since signaling peptides that share common ancestry often also exert similar functions, their study can provide fundamental insight into the common biology of species that span diverse phyla, e.g., from protostome model organisms, such as the fruit fly, the nematode *Caenorhabditis*, and the sea hare *Aplysia*, to deuterostomes, including zebrafish and humans. Examples of ancient signaling peptides include those of the insulin family, neuropeptide Y peptides, and oxytocin/vasopressin-like peptides (Elphick et al. 2018).

While important to know, the evolutionary origin of signaling peptides can be challenging to uncover. Because of their small size, signaling peptides often lack sufficient sequence conservation to establish homology, especially if peptides are compared between distantly related species. Additionally, many signaling peptide precursors do not contain well-defined structural or functional domains that could help in determining their relatedness. Repetitive sequence regions of varying frequency and lengths are also common, further hampering efforts to establish relatedness between signaling peptides from divergent species. Consequently, many homologous signaling peptides are diversified to a point beyond recognition, even though they share common ancestry.

There is alternative evidence that can be gathered to establish homology. For example, conserved structures of the peptide-encoding genes (the number, positions, and phases of introns) can in some cases be used to determine homology (Mair et al. 2000; Yañez-Guerra et al. 2020; Zhang et al. 2020). Furthermore, comparative sequence analysis of the target receptor, which are often G protein-coupled receptors (GPCRs), can assist in establishing orthology. Because of their larger size and well-defined sequence structures, compared to their peptide ligands, GPCRs are more amendable to phylogenetic analyses. GPCR receptor-ligand pairs are typically stably associated throughout evolution. Thus, orthology between divergent signaling peptide ligands may be inferred from orthologies of their receptors (Mirabeau and Joly 2013). This approach has been successfully used to find common ancestry of, for example, deuterostome gonadotropin-releasing hormone and protostome adipokinetic hormone, even though the peptide ligands themselves only share a single amino acid (Grimmelikhuijzen and Hauser 2012). Although these methods have proven successful in tracing the evolution of some signaling peptides, the origin and relatedness of many important receptor-ligand systems remains unknown.

In this paper, we use a novel approach for inferring homology based on a group of venom peptides that share sequence and functional similarity with somatostatin and its related peptides (SSRPs), a family of hormones that was until recently believed to be restricted to chordates. As we show here, the venom peptides unravel the existence of an SSRP-like signaling system in protostomes and track the evolution of this system to early Bilateria.

The venom peptides used here are produced by a highly-biodiverse marine lineage, the predatory cone snails (*Conus*). Cone snail venoms are extremely complex, with hundreds of different components per species, which can be grouped into gene families (Woodward et al. 1990). Each venom peptide family has a conserved signal sequence, but the mature secreted peptide is subject to accelerated evolution, resulting in the venom of each species having its own distinctive complement of peptides (Li et al. 2017). Endogenous signaling peptides, widely distributed across all molluscs, can be recruited for expression in cone snail venom – this has previously been demonstrated for oxytocin (Cruz et al. 1987) and insulin (Safavi-Hemami et al. 2016). In a recent paper, we identified another family of signaling peptide-like toxins in *Conus* venom with significant sequence similarity to somatostatin (Ramiro et al. 2021). Whether these somatostatin-like cone snail toxins (consomatins) evolved from an endogenous somatostatin-like signaling peptide was not addressed.

Somatostatin (SS) is a highly conserved chordate hormone, which was initially discovered as an inhibitor of growth hormone release from the hypothalamus (Brazeau et al. 1973). It has since been found to serve diverse physiological functions, including the inhibition of insulin and glucagon release from the pancreas, regulation of gut motility, and as an analgesic, anxiolytic and anticancer agent (Martinez 2015). SS was the first discovered member of a multigene family of peptides that all evolved from an ancestral SS-like gene. During chordate evolution the ancestral SS gene duplicated in multiple rounds, such that there are four peptides related to SS in mammals: somatostatin (SS), cortistatin (CST), urotensin-II (UII) and urotensin-II related peptide (URP), and 8 such genes in teleost fish (**Figure 1A**) (Tostivint et al. 2014). SS and CST are inhibitors of hormone secretion, inflammation, and pain while UII and URP are important regulators of cardiovascular activity (Moller et al. 2003; Vaudry et al. 2010). Several studies have also suggested partially overlapping effects of these peptides. For example, UII is primarily known as a potent vasoconstrictor, but it also shares several functions with SS, such as inhibition of insulin secretion and regulation of food intake (Ong et al. 2008). In this article, we collectively refer to these peptides as somatostatin-related peptides (SSRPs).

**Figure 1.**
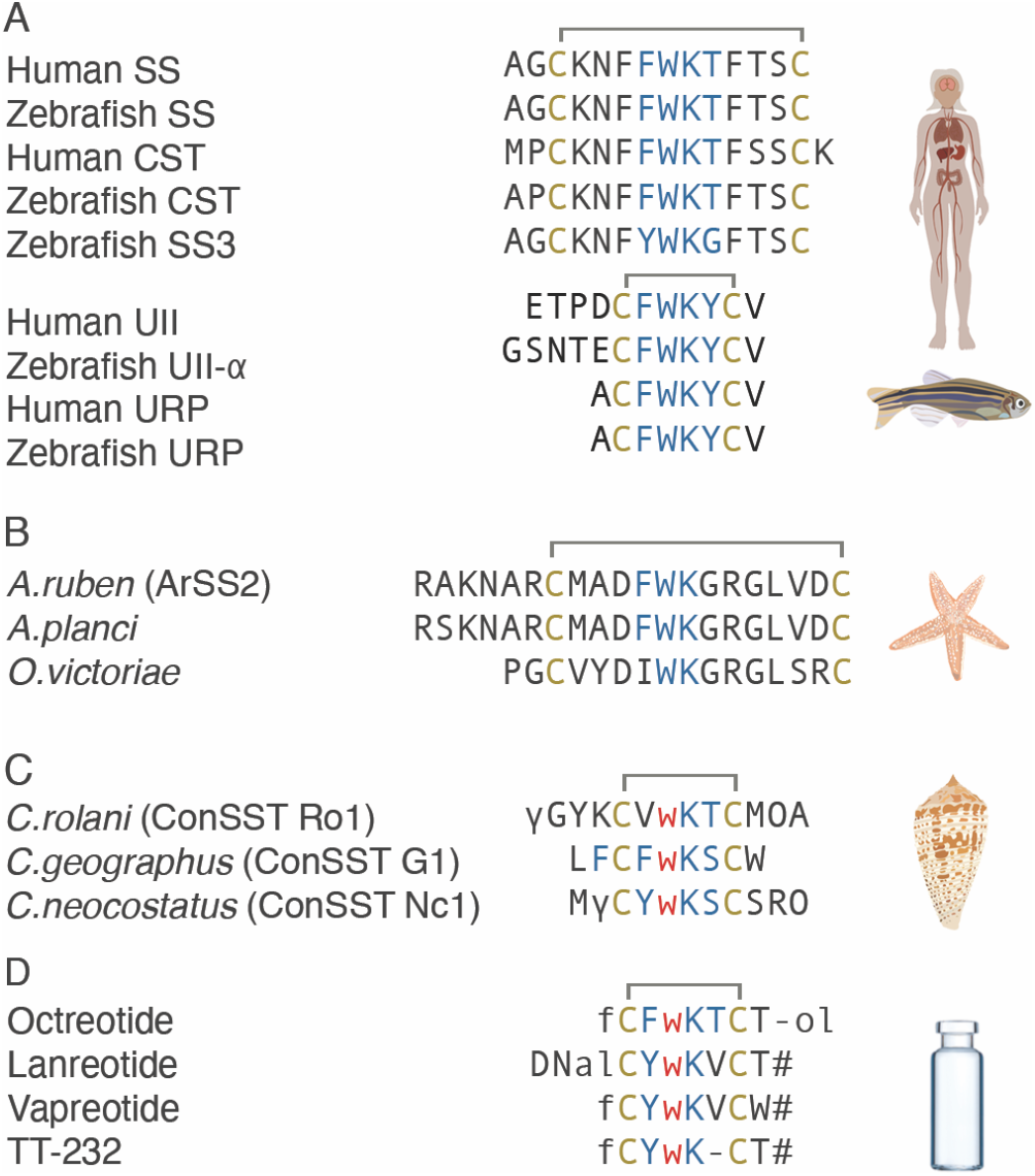
Somatostatin and its related peptides (SSRP) from (A) vertebrates (human and zebrafish), and (B) echinoderms, (C) consomatins from fish-hunting cone snails, and (D) therapeutic somatostatin (SS) analogs. Cysteines and amino acids of the core SS receptor binding motif are shown in yellow and blue, respectively. Modifications: γ=γ-carboxyglutamate; w=D-tryptophan (shown in red); O=hydroxyproline, f=D-phenylalanine; #=amidation; -ol=alcohol, DNal=3-(2-Naphthyl)-L-alanine

SSRPs exert their function through activating various members of the GPCR family. In humans, both SS and CST activate all of the five SS receptors, SST_1-5_ (Moller et al. 2003). In addition, CST also activates the ghrelin receptor and Mas-Related G Protein-Coupled Receptor-X2 (MRGPRX2) (Deghenghi et al. 2001; Robas et al. 2003). UII and URP both bind to and activate the urotensin receptor (UT) (Vaudry et al. 2010). As for their ligands, SST_1-5_ and UT are thought to have originated from a common ancestor during chordate evolution (Tostivint et al. 2014). SSRPs were initially believed to be exclusively found in chordates, but the discovery of an SS-like peptide in the starfish *Asterias rubens* (phylum Echinodermata) capable of activating an SST-like receptor established the presence of an SSRP signaling system in deuterostome invertebrates and suggested a more ancient evolutionary history of SSRPs (**Figure 1B**) (Semmens et al. 2016; Zhang et al. 2020). Furthermore, our recent discovery of consomatins in the venom of fish-hunting cone snails that selectively activate different subtypes of the human SST1-5 suggested the existence of an SSRP-like signaling system in protostomes that may have given rise to the consomatin family (**Figure 1C**) (Ramiro et al. 2021).

Here, by tracing the evolution of consomatins we unravel the existence of homologous peptides in diverse protostome phyla that, based on several lines of evidence, are proposed to be orthologs of chordate SSRPs. Specifically, we find that consomatins are derived from SSRP-like signaling peptides that are expressed in neuroendocrine tissue of mollusks, annelids, and several other protostome phyla. The signaling SSRP-like gene experiences purifying selection to target a conserved endogenous receptor, but once duplicated and recruited into venom, rapidly diversifies to target receptors in a shifting mix of prey and, potentially, predators and competitors. Collectively, our findings shed light on the broad existence and early evolution of the SSRP-like signaling system in Bilateria and provide a new paradigm for the use of animal toxins to track the evolution of important signaling systems across diverse prey taxa.

## Results

### Evolution of diverse cone snail toxins with similarity to chordate SSRPs

Recently, we identified and characterized a novel family of somatostatin-like toxins (consomatins) in fish-hunting cone snails of the *Asprella* and *Gastridium* clade that target the SS signaling system in vertebrates (Ramiro et al. 2021). Consomatins share several conserved residues with chordate SSs, selectively activate different subtype of the human SS receptor, and display a minimized disulfide scaffold and a D-amino acid that is also found in synthetic SS drug analogs, such as Octreotide and Lanreotide (**Figure 1C-D**). In our previous study, we specifically searched for toxins that share at least three of the four residues important for activating the human SS receptor, or have conservative substitutions in these positions (residues 7-10: FWKT, numbering based on human SS-14). Here, to elucidate the full diversity of consomatins that not only share sequence similarity with SS but also other members of the SSRP family, we more broadly searched for sequences that display the WK motif (residues 8-9) and disulfide loop characteristic for these peptides (see **Figure 1A**).

By searching the genomes and venom gland transcriptomes of a phylogenetically diverse set of 25 cone snail species, we identified 52 consomatin sequences containing these features (**Figures 2A and S1**, see **Supporting File 1 and 2** for sequences and accession numbers of all datasets interrogated in this study). All sequences belong to the C toxin gene superfamily that is defined by a conserved N-terminal signal sequence involved in translocation to the endoplasmic reticulum and secretory pathway.

**Figure 2.**
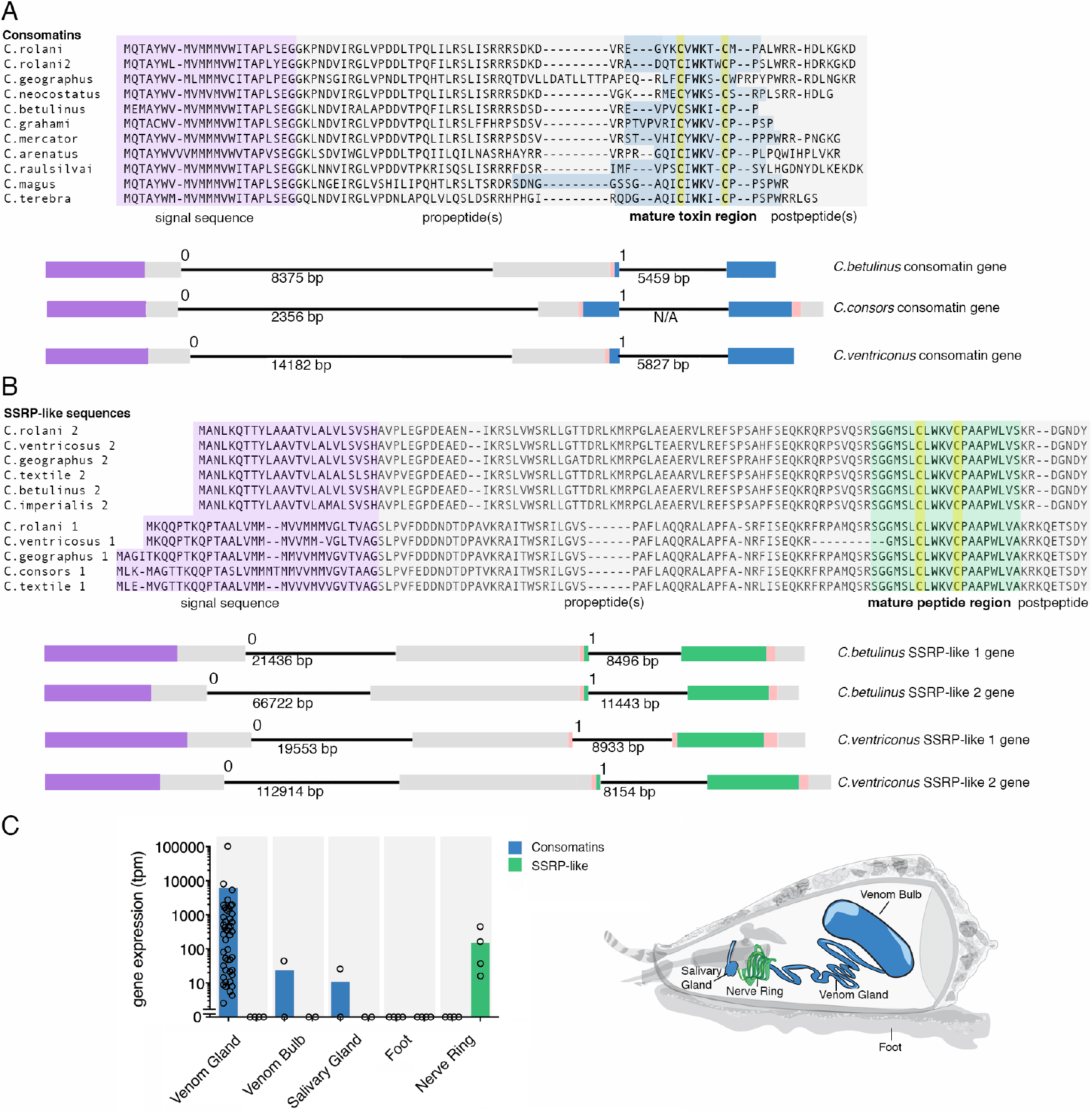
Consomatins and signaling SSRP-like peptides are homologs with contrasting expression patterns. Comparative sequence alignments and gene structures of selected (**A**) consomatins and (**B**) signaling SSRP-like sequences retrieved from transcriptome and genome data. The position of the signal sequence (purple), propeptides (gray), mature toxin region (blue), mature signaling SSRP-like peptide region (green), and postpeptides (gray) are shown below alignments. Gene structures are shown using the same color codes. (**C**) Mean gene expression levels of consomatins and signaling SSRP-like sequences in the venom gland (n=46), venom bulb (n=2), salivary gland (n=2), foot (n=4), and circumoesophageal nerve ring (n=4).

Consomatin genes contain 3 exons (**Figure 2A**): the first exon covers the signal sequence and N-terminal part of the pro-region, the second exon encodes the C-terminal part of the pro-region, as well as the N-terminal processing site and first 1-5 amino acids of the mature peptide, and the last exon spans the remaining part of the mature peptide, and in most cases a C-terminal processing site and the remaining post-peptide region. The two introns are of phase 0 and 1, respectively, with the first intron being longer.

### Consomatins evolved from an ancestral SSRP-like precursor gene

The high sequence similarity of consomatins with chordate SSRPs suggested that these toxins may have originated from a molluscan SSRP-like gene involved in neuroendocrine signaling. To investigate this possibility, we sequenced the transcriptomes of the circumoesophageal nerve ring, an arrangement of nerve ganglia known to secrete neuroendocrine signaling peptides (Safavi-Hemami et al. 2016) from four divergent cone-snail species, *Conus rolani, Conus geographus, Conus textile,* and *Conus imperialis*. Homology searching for an SSRP-like gene led to the identification of two highly expressed sequences encoding prepropeptides that share the WK motif and disulfide loop with consomatins and, therefore also with vertebrate SSRPs (**Figure 2B, Supporting Files 1 and 3**). The two SSRP-like genes (SSRP-like 1 and SSRP-like 2) have common characteristics of peptide preprohormones (N-terminal signal sequence and canonical proteolytic processing sites), and are expressed specifically in the nerve ring, whereas consomatin sequences are expressed primarily in the venom gland, and at much lower levels in the venom bulb, a muscular extension of the gland that plays a role in venom delivery, and the salivary gland, an accessory gland located in close proximity to the venom gland (**Figure 2C**). To specifically determine if the consomatin and the SSRP-like signaling genes evolved from a common, ancestral SSRP-like predecessor, we interrogated the available genomes of *Conus betulinus, Conus consors,* and *Conus ventricosus*. We found that the organization of the signaling genes mirrors that of the venom genes, including the number, site, and phases of the introns, and the location of the mature peptides within the genes (**Figure 2A and B**). These highly conserved structural features corroborate that consomatins evolved by gene duplication of a signaling SSRP-like gene expressed in the snail’s neuroendocrine system. Whether consomatins emerged from only one or both SSRP-like signaling genes could not be determined. However, as evident by phylogenetic analysis, following the duplication event(s), consomatin genes neofunctionalized and proliferated leading to a great diversification of this venom peptide family (**Figure 3A**).

**Figure 3.**
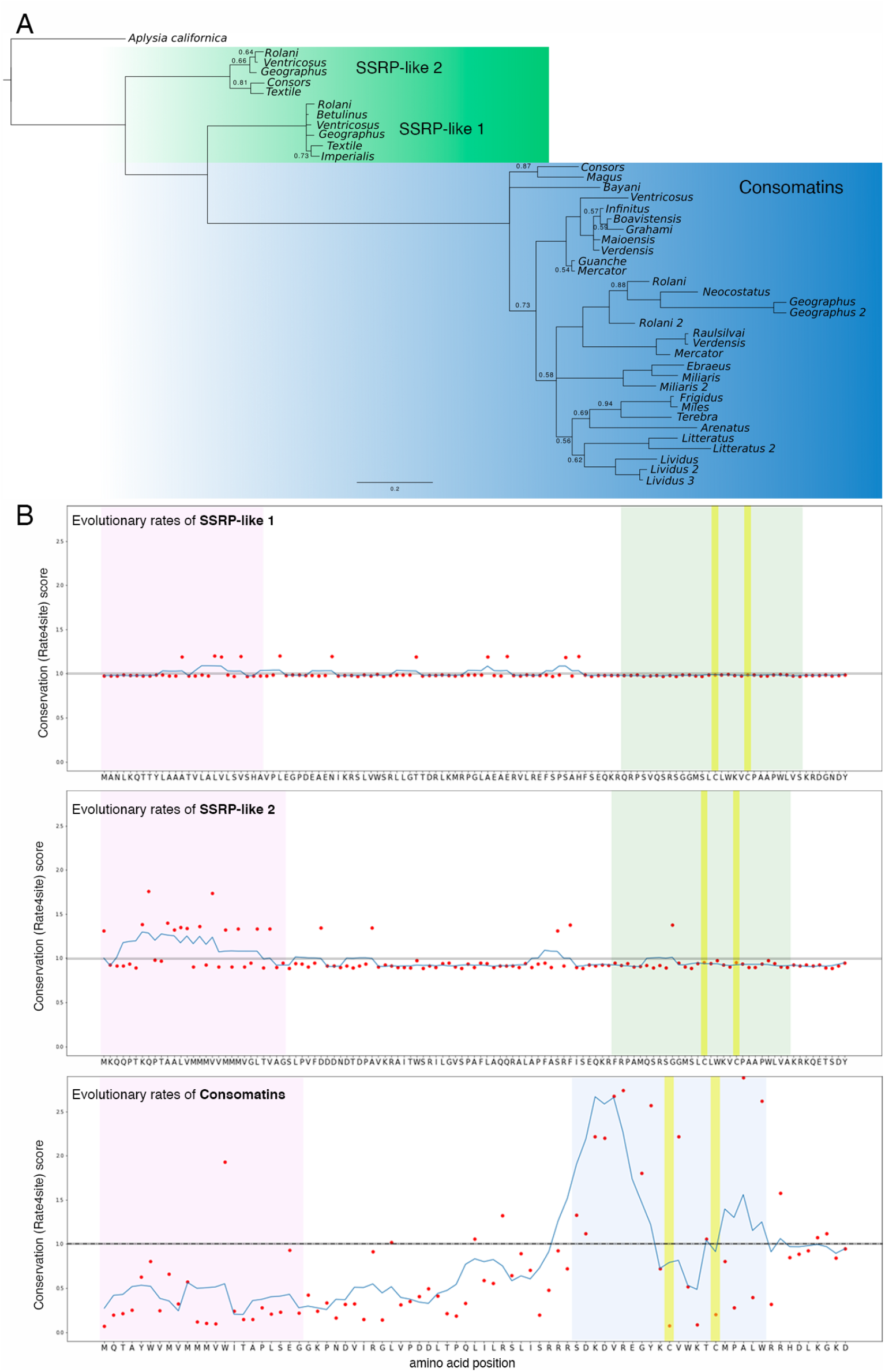
Consomatins and signaling SSRP-like sequences show contrasting evolutionary patterns. (**A**) Bayesian gene tree of consomatin and nerve ring SSRP-like sequences from diverse cone snail species. Branch lengths are proportional to estimated amino-acid change, highlighting the speed with which consomatins have diverged, as compared to the nerve ring sequences. Only posterior probabilities values below 0.95 are depicted for visual clarity. (**B**) Position-specific evolutionary rates of SSRP-like 1 (top), SSRP-like 2 (middle), and consomatin (bottom) sequences as determined using rate4site. Amino acids of the corresponding sequences from *Conus rolani* are shown for clarity. Color codes are the same as in Figure 2.

### Contrasting evolutionary patterns between consomatins and signaling SSRP-like genes

In contrast to the hyper-diverse consomatin family, the two signaling SSRP-like genes are almost identical to one another within and between the four different cone snail species (mean amino-acid sequence identity of 50.3 % between the genes and 96.6 % and 91.5 % between species for each of the two genes). This is further highlighted by their close grouping and short branches in phylogenetic analyses (**Figure 3A**). To determine whether the signaling genes have been subject to negative/purifying selection, we estimated the pairwise dN/dS ratios between genes from different cone snails (**Supporting File 4**). The dN/dS ratio is the rate of nonsynonymous to synonymous mutations, where ratio above 1 indicates positive selection, while a ratio below 1 indicates purifying selection. Based on estimates of pairwise dN/dS ratios, the signaling genes appear to have been subject to negative/purifying selection (SSRP-like gene 1: mean dN/dS = 0.34, range: 0.08-1.20; SSRP-like gene 2: mean dN/dS= 0.57, range: 0.11-1.02). This is also confirmed by comparing M7 and M8 models in the program PAML, which estimates the likelihood that some sites evolved under positive selection. This analysis showed that none of the sites appear to diverge from negative or neutral selection in any of the two genes (**Table 1**).

**Table 1.**
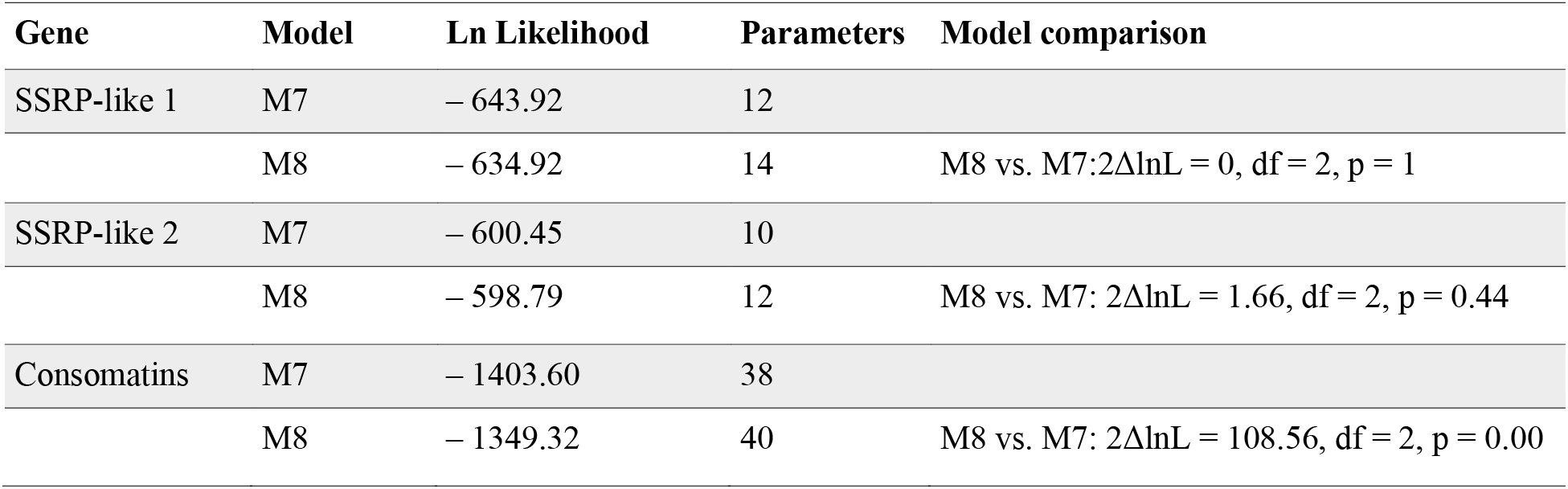
Models and statistical values of pairwise dN/dS ratios between gene families (SSRP-like 1, SSRP-like 2, and consomatins) from different cone snails.

To estimate the evolutionary rate of the consomatin genes we randomly selected 20 transcriptome sequences. Pairwise comparison gave average dN/dS = 3.04 with 88 % of comparisons above 1 (**Supporting File 4**). Tests for positive selection (M7 vs M8) strongly favors a model, where 26 % of the 62 aligned codons have evolved with an average dN/dS-ratio of 6.94, in contrast to a model constrained to negative or neutral evolution at all sites (model statistics can be seen in **Table 1**). Positive selection is particularly evident at sites around the mature toxins, except for the cysteine residues that form the disulfide loop and the WK-motif, which are all highly conserved.

Plotting the conservation (Rate4site) scores for consomatins can further be used to visualize how the different regions of the toxins precursor evolve at different rates. While the signal sequence and pro-regions are all well conserved, amino acids within the mature toxins are highly divergent, apart from the two cysteines and the WK motif. In stark contrast, the signaling SSRP-like sequences are highly conserved all throughout the preprohormone (**Figure 3B-C**).

### The signaling SSRP-like genes are ubiquitously found in all classes of mollusca

We hypothesized that the two SSRP-like genes found in the circumoesophageal nerve ring of cone snails encode secreted peptides involved in neuroendocrine signaling. Since signaling peptides are generally well conserved within phyla, we searched for similar genes in other mollusks. This led to identification of highly similar genes in all molluscan classes (Gastropoda, Bivalvia, Polyplacophora, Cephalopoda, Scaphopoda, Aplacophora and Monoplacophora, see **Supporting File 5** for sequences). In particular, we found that the cone snail SSRP-like genes are orthologous to a previously described gene from the sea hare *Aplysia californica* that encodes a secreted, bioactive peptide expressed in various ganglia (buccal, cerebral, pleural-pedal and abdominal) (Romanova et al. 2012). Based on its sequence similarity to vertebrate UII this peptide was named *Aplysia* UII (apUII). ApUII shares 40 % identity with the predicted cone snail signaling SSRP-like peptide. This finding confirms that the SSRP-like gene family that gave rise to consomatins encodes a molluscan neuroendocrine peptide that shares sequence similarity with members of the chordate SSRP family. While the *A. californica* genome assembly was not of sufficient quality to interrogate the structure of the apUII-encoding gene, analysis of several other molluscan genomes, including that of the golden apple snail, *Pomacea canaliculata,* confirmed the same exon/intron structure as identified for the consomatin and the cone snail signaling SSRP-like genes (**Figure S2A**).

### Consomatins expressed in worm-hunting species unravel the existence of an orthologous signaling system in annelids

Previously, we showed that two consomatins from fish-hunting cone snails of the *Asprella* and *Gastridium* clade share several important amino acids with vertebrate SSs and selectively activate the human SST_1,4_ and SST_2_ with low micromolar and low nanomolar potency, respectively (Ramiro et al. 2021). We further noticed that consomatins were also expressed in some cone snail species that prey on annelid worms which strongly suggested the existence of an SSRP-like signaling system in annelids. Here, by searching the genomes and transcriptomes of 19 species of worm-hunting cone snails we first confirm the wide distribution of consomatins in various clades of worm hunters (48 sequences from 19 species belonging to 10 clades). We then proceeded to use the molluscan signaling SSRP-like sequences to query similar genes in a wide range of annelid genomes and transcriptomes. By doing so, we find 19 SSRP-like sequences from 14 species belonging to both classes of Annelida (Polychaeta and Clitellata), demonstrating the wide distribution and expression of SSRP-like genes in this large protostome phylum (**Supporting File 6**). Notably, SSRP-like sequences are not restricted to marine species but are, for example, also present in the earthworm *Lumbricus rubella*.

Several, but not all, annelid SSRP-like genes have gene structures that are identical to the molluscan counterpart: 3 exons that are separated by a phase 0 and phase 1 intron with the signal sequence located on exon 1, and with the mature peptide starting at the 3’ end of exon 2 and extending into exon 3, where the conserved WK-motif and disulfide loop is located (**Figure S2**). This gene structure and high similarity to the molluscan genes strongly suggests that these systems are orthologous. However, not all annelids strictly adhere to this canonical SSRP-like gene structure. In *Dimorphilus gyrociliatus,* an annelid with a miniaturized genome (Martín-Durán et al. 2021), two of the three SSRP-like genes lack the second intron, while in *Streblospio benedicti*, an annelid with an unusually large genome (Zakas et al. 2021), a third intron interrupting the signal sequence can be found (**Figure S2**).

Our findings on the existence of SSRP-like sequences in diverse classes of Mollusca and Annelida prompted us to look for similar sequences in other metazoan phyla. This search led to the identification of SSRP-like sequences in three additional phyla: Brachiopoda, Rotifera and Xenacoelomorpha (**Supporting File 7**, and where available their conserved genes, **Figure S2**). The phylogenetic position of xenacoelomorphs is still debated, but the current consensus is that xenacoelomorhps are bilaterians that diverged before the major Protostomia-Deuterostomia split (Cannon et al. 2016; Hejnol and Pang 2016; Jondelius et al. 2019; Kapli and Telford 2020). Notably, SSRP-like peptides appear to be absent from ecdyzozoans, a group of protostomes that includes nematode worms and arthropods. Thus, SSRP-like sequences are found across many but not all branches of bilaterians (**Figure 4**).

**Figure 4.**
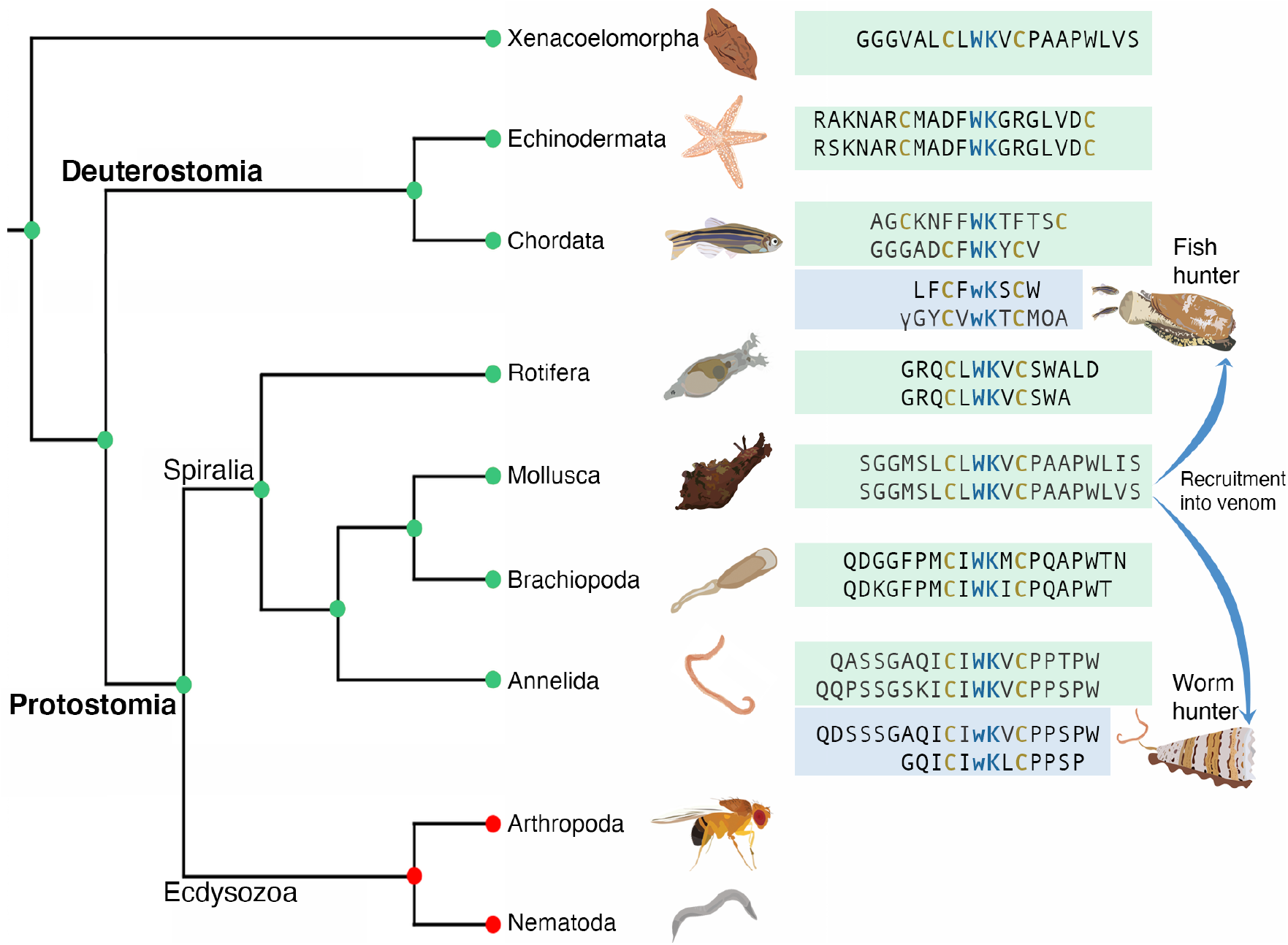
Distribution and selected sequences of SSRP-like genes in diverse bilaterian phyla. Cladogram showing proposed retention (green circles) and loss (red circles) of the SSRP-like gene across diverse animal phyla. Green boxes show select sequences of signaling SSRP-like genes, blue boxes depict highly similar consomatin sequences identified in fish- and worm-hunting species of cone snails. Sequences (from top to bottom): *Xenoturbella bocki*, *Asterias rubens* (ArSS2), *Acanthaster planci* (Apla_2), *Homo sapiens* (SS, identical sequence present in *Danio rerio*), *Danio rerio* (UII), *Conus geographus* (consomatin G1), *Conus rolani* (consomatin Ro1), *Rotaria sordida*, *Rotaria tardigrada*, *Aplysia californica* (apUII), *Conus textile*, *Linguna anatina*, *Linguna anatina* 2, *Capitella teleta* (SSRP-like peptide 1), *Platynereis dumerilii*, *Conus floridulus* (consomatin Fr1), *Conus tribblei* (consomatin Tl1). Modifications: γ=γ-carboxyglutamate; w=D-tryptophan; O=hydroxyproline.

### Protostome SSRP-like peptides activate receptors that share sequence similarity with chordate somatostatin receptors

To further investigate the function of protostome SSRP-like sequences, we synthesized the predicted mature peptides of Ct-SSRPL1 from the genome and whole-body transcriptome of the annelid *Capitella teleta* and Cr-SSRPL1 from the nerve ring transcriptome of *Conus rolani* and tested them at their predicted receptor pairs; an annelid candidate receptor from the transcriptome of *C. teleta* (Ct-SSTL1), and a molluscan candidate receptor from the *Conus betulinus* genomic DNA assembly (Cb-SSTL1). These orphan receptors grouped with the human SSTs in maximum likelihood tree analyses, suggesting that they are members of an invertebrate SSRP signaling system (**Figure 5A and S3**). We note that the molluscan receptors with the highest sequence similarity to the chordate SS receptors are allatostatin C (ASTC) receptors (**Figure 5A**). ASTCs are protostome peptides with some resemblance of SS (a C-terminally located cyclic peptide but no WK motif) and are believed to be related to SS based on the similarity of their receptors to the SS receptor and their inhibitory functions (Mirabeau and Joly 2013; Veenstra 2009). ASTC is also present in mollusks (Zhang et al. 2018), but while we could identify ASTC in the cone snail nerve ring transcriptomes (data not shown) the ASTC ligand-receptor system was not further investigated here.

**Figure 5.**
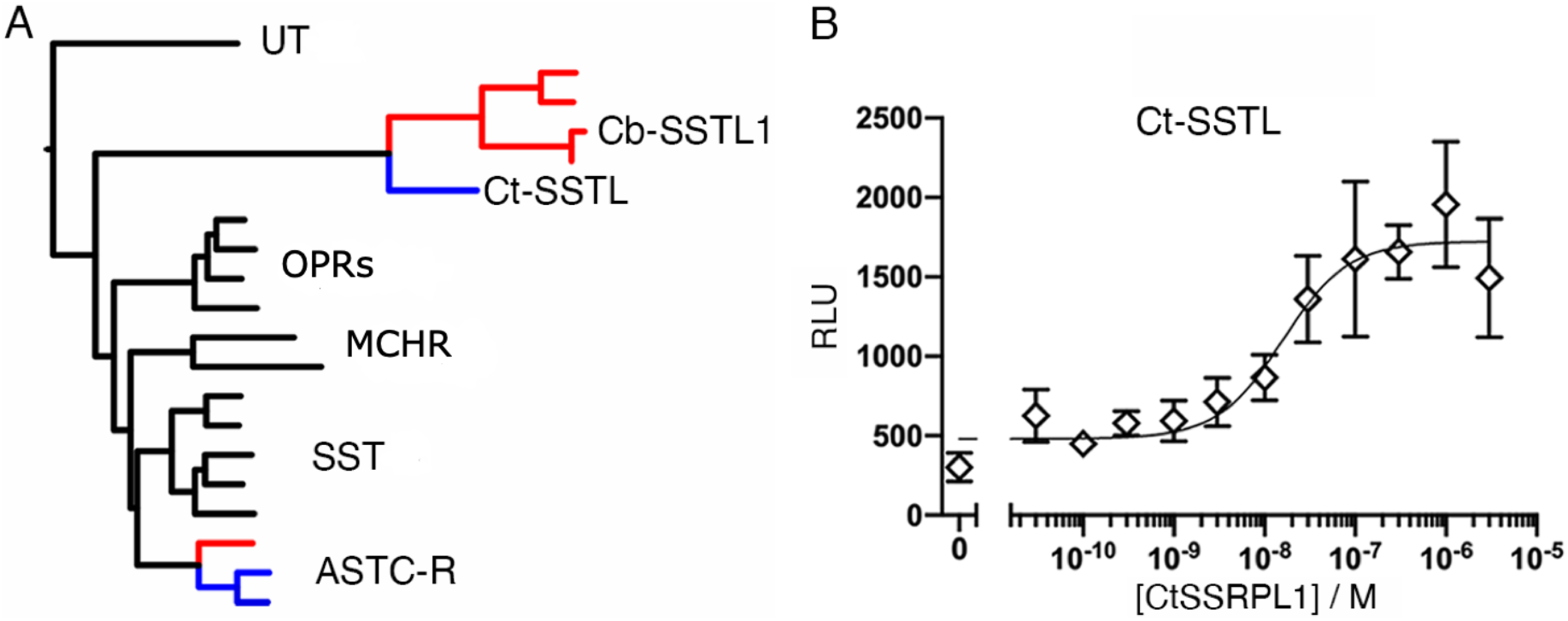
Ct-SSL1 clusters with human SSRP and related receptors and is activated by Ct-SSRP1. (**A**) Maximum likelihood tree analysis of human SSRP receptors (SST and UT) and closely related receptors (Opioid receptors (OPRs) and melanin-concentrating hormone receptors (MCHR)) and the ASTC-R from the annelid *Platynereis dumerilli* with orphan receptors from *Conus betulinus* (red branches) and *Capitella teleta* (blue branches). The tree was rooted with human galanin and kisspeptin receptors (not shown). The location of Ct-SSTL1 and Cb-SSTL1 are shown. The full tree with bootstrap values and labels can be found in **Figure S3**. (**B**) Concentration-response curve of Ct-SSRPL1 at the Ct-SSTL1 receptor in the PRESTO-Tango assay (error bars represent SD). The experiment was performed in three independent repeats, and the mean EC_50_ value was 13.1 nM (pEC50 ± CI95 = 7.88±0.65).

The *C. teleta* peptide, Ct-SSRPL1, showed strong activation of its predicted receptor, Ct-SSTL1, with an EC_50_ of 13.1 nM (pEC50±CI95 = 7.88±0.65) (**Figure 5B**). This finding suggests that the Ct-SSTL1 receptor is indeed the functional target of this peptide. In contrast, the cone snail peptide, Cr-SSRPL1, showed no robust response at its predicted receptor, indicating that this receptor may not represent the biological target of this peptide or that the receptor was not expressed in its functional form (data not shown). However, when applied to the Ct-SSTL1 receptor at 30 μM, the highest concentration tested, Cr-SSRPL1 showed approximately 50 % activation, and was also able to activate the human SST_2_ with micromolar potency (**Figure S5**).

### Consomatins evolve to target the SSRP-like signaling system in prey

Collectively, our findings suggest that early in the evolution of cone snails, a duplicated SSRP-like gene that functions in neuroendocrine signaling in many bilaterians was recruited for expression in the venom gland. As cone snails radiated and diversified ecologically, the venom gene greatly proliferated and diverged in sequence.

The prey of ancestral cone snails were polychaete worms (Duda et al. 2001; Puillandre et al. 2014). Some cone-snail lineages later shifted to prey on mollusks, and others on fish (Kohn 1956; Olivera et al. 2015; Puillandre et al. 2014). Having established the existence of SSRP-like sequences in these diverse prey taxa allowed us to test if the great diversification of consomatins occurs in response to biotic interactions with different prey. In other words, if consomatins target the SSRP-like receptor of prey species, then they would be expected to resemble the native (signaling) SSRP-like peptides of those prey species. To investigate this hypothesis, we performed principal component analysis (PCA) of vertebrate SSRPs (including fish and human sequences), signaling SSRP-like sequences from mollusks and annelids as well as a large set of consomatins. PCA transforms a large group of correlated variables into a smaller set of uncorrelated variables (the principal components) that are more easily interpretable. Here, the scored variables included length, aromaticity, molecular weight, isoelectric point, and count of the 150 most common 1-3mers (see Material and Methods). Because we established that the mature region of consomatins is almost exclusively located on a single exon (exon 3) we were able to utilize recent exon capture data of this exon from 247 distinct species of cone snails representing all major branches of the *Conus* phylogenetic tree (Phuong et al. 2019) (**see Table S1** for all species and accession numbers used in this study). Using these datasets, we identified 502 consomatin genes of which 71 and 431 were from 30 fish-hunting and 131 worm-hunting species, respectively (**Supporting File 8**). Consistent with transcriptome analysis, consomatin genes greatly proliferated in some lineages (e.g., *Africonus, Virroconus, Dauciconus, and Lividoconus*), and were seemingly lost in others (e.g., *Textilia* and *Tesseliconus*) (**Figure 6A**). Interestingly, consomatin genes could not be detected in any of the 22 snail-hunting species analyzed. This may point to a loss of this gene during the shift from worm- to snail-hunting behavior which is believed to have only occurred once (Duda et al. 2001).

**Figure 6.**
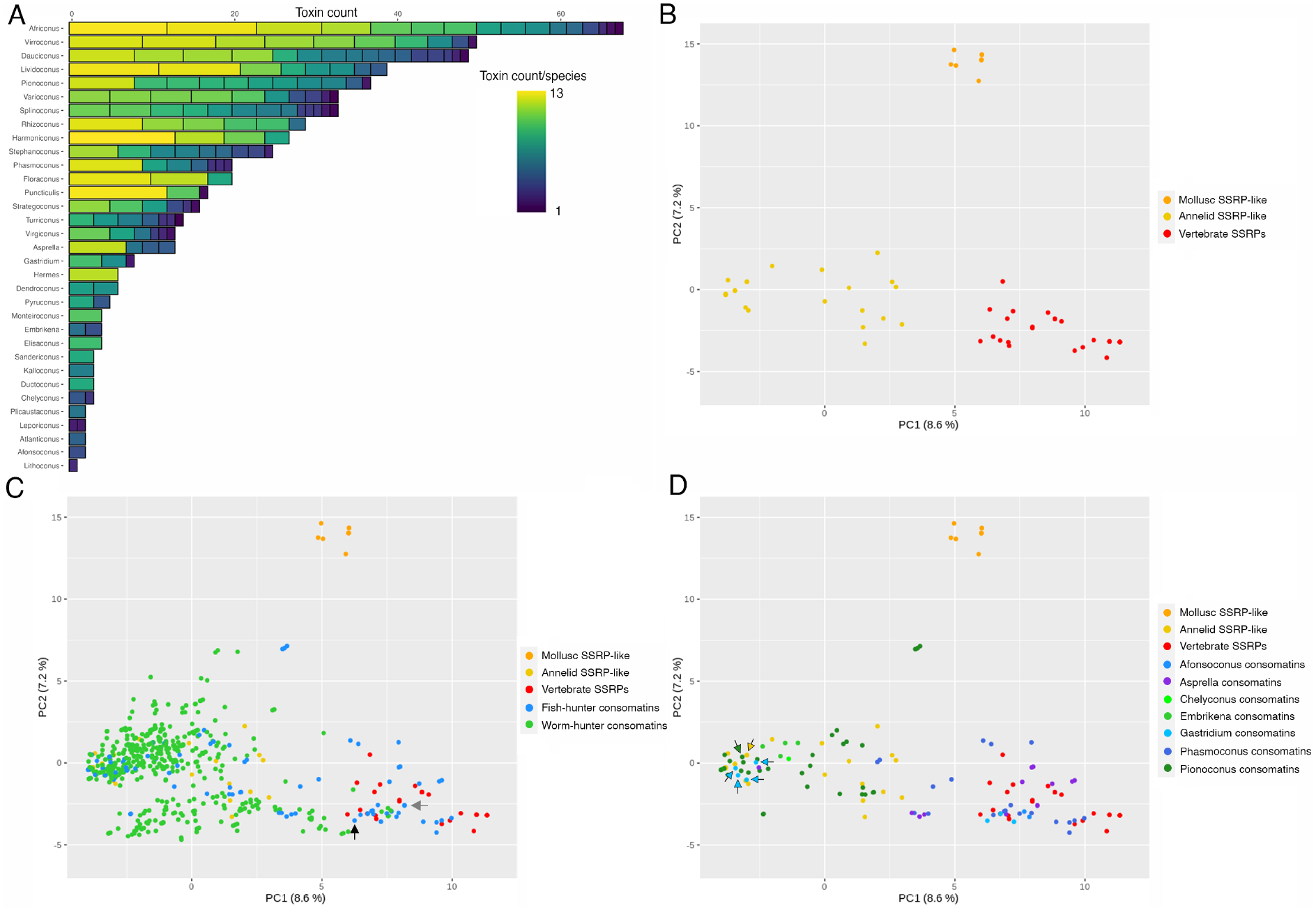
Diversity of consomatins and their correlation with prey peptides. (**A**) Bar graph of the consomatin count in the different cone snail lineages investigated in this study. Each bar is comprised of smaller elements representing the individual species of the lineage. Both the length and color of smaller bars represents the number of consomatins. Consomatins could not be identified in the following lineages: *Fraterconus*, *Klemaeconus*, *Lindaconus*, *Textilia*, *Tesselliconus*, and all 5 lineages of snail hunters included here: *Eugeniconus*, *Conus*, *Leptoconus*, *Darioconus* and *Cylinder*. (**B**) Principal component analysis (PCA) of characteristics of vertebrate SSRPs, annelid and molluscan SSRP-like sequences. (**C**) Same analysis as in (**B**) including consomatins from fish-and worm-hunting cone snails. (**D**) PCA showing SSRPs and SSRP-like signaling peptides from vertebrates, annelids, and mollusks as well as consomatins from fish-hunters separated into clades.

In analyses of the signaling SSRP-like gene alone, the first two principal components separate sequences from mollusks, annelids, and vertebrates into distinct clusters (**Figure 6B**). In analyses of the signaling SSRP-like peptides together with the consomatin sequences, consomatins noticeably group away from the molluscan signaling genes they derived from, andx towards the annelid and vertebrate sequences (**Figure 6C**). This pattern is consistent with the hypothesis that consomatins diverged to target SSRP-like receptors in prey. Indeed, consomatin sequences from fish hunters, that were previously shown to selectively target different subtypes of the human SST receptors, closely group with vertebrate SSRPs, including those from fish and human (see black and gray arrows for consomatin G1 and Ro1, respectively, **Figure 6C**). In contrast, consomatins identified from worm hunters cluster almost exclusively towards the annelid SSRP-like sequences, again suggesting that consomatins evolve to target the SSRP-like signaling system of specific prey. Notably, a small number of consomatins from the worm-hunter, *Conus glaucus*, and several worm-hunting species of the *Africonus* clade are placed in the vertebrate region of the plot potentially indicating that these species may also prey on fish (as observed for some other worm hunters (Aman et al. 2015)) or may use these toxins for defensive purposes.

Except for these sequences, all other consomatins that group with vertebrate SSRPs are from fish-hunting species. This not only includes consomatins from *C. rolani* (*Asprella* clade) and *C. geographus* (*Gastridium* clade) known to activate the human SS receptors (Ramiro et al. 2021), but also sequences retrieved from other species belonging to these two clades (**Figure 6D**). Additionally, sequences from two other phylogenetically distinct clades of fish hunters, *Phasmoconus* and *Afonsoconus*, also group closely with vertebrate SSRPs. This includes the sequence of a recently identified consomatin from *Conus ochroleucus* (*Phasmoconus*) that is nearly identical to the SS drug analog octreotide and potently activates the human SST_1-4_ (Acaytan et al., unpublished). Thus, PCA analysis and comparative sequence analysis highlight how some, but not all fish hunters adapted the duplicated signaling SSRP-like gene to specifically target the receptors of their fish prey.

Notably, a large number of consomatins from fish hunters do not cluster with chordate SSRPs but with those from annelids. Most of these sequences belong to a single clade, *Pionoconus*, that comprises some of the best studied cone snail species. Interestingly, this clade includes *Conus magus*, a species that is known to prey on annelids as a juvenile (Nybakken and Perron 1988), indicating that its consomatin sequence may have adapted to target the SSRP-like receptor in annelid prey, and possibly, that other members of this clade also prey on annelids as juveniles. Grouping of the *C. magus* peptide with SSRP-like sequences identified in annelids that are closely related to the prey of juvenile *C. magus* strongly supports this hypothesis (see green and yellow arrows in **Figure 6D** representing the *C. magus* and the annelid *C. teleta* sequence, respectively).

Finally, we note that the position of consomatins from fish hunters in the PCA plot also appears to correlate with predation strategy. All members of the *Pionoconus* clade are all taser-and-tether hunters, a predation strategy characterized by a rapid immobilization of prey facilitated by the action of toxins that rapidly modulate ion channels of the nervous and locomotor systems (Olivera et al. 2015). In contrast, species of the *Asprella* clade, and potentially also the *Phasmoconus* clade, are ambush-and-assess hunters, a hunting strategy that is characterized by a very slow onset of action of their toxins (Ramiro et al. 2021). The third predation strategy, net-hunting, in which toxins are released into the water, has only been observed in *Gastridium* clade and suggested to be used by some *Afonsoconus* species (Ahorukomeye et al. 2019). The PCA analysis reveals a clear pattern: sequences derived from taser-and-tether hunters group with annelid SSRP-like sequences, whereas toxins from net or ambush-and-assess hunters group with vertebrate SSRPs (**Figure 6D**). PCA grouping according to predation strategy, rather than phylogeny alone, is best illustrated by two closely related species of the *Gastridium* clade: consomatins from *Conus obscurus*, a taser-and-tether hunter (Olivera et al. 2015), closely groups with the annelid sequences (light blue arrows in Figure 6D) whereas consomatins from the net-hunter *C. geographus* group with vertebrate SSRPs. Based on this pattern it can be hypothesized that many, if not all, taser-and-tether hunters prey on annelids in their early developmental stages.

## Discussion

Our investigations into the evolutionary origin of consomatins exposed the existence of an SSRP-like signaling system in diverse protostome phyla that gave rise to the consomatin family. Several lines of evidence suggest that this signaling system shares a common origin with chordate SSRPs, a family of peptide hormones that includes SS, CST, UII, and URP, and their respective receptors, which in humans is the SST_1-5_ and the UT.

First, all protostome sequences discovered here share the conserved WK motif known to be of critical importance for SST1-5 and UT activation (**Figure 4**) (Moller et al. 2003). We note that the residue preceding this motif in deuterostomes is almost always aromatic (Phe or Tyr) but aliphatic in protostomes (Ile or Leu) pointing towards important differences in the receptor binding mode of the protostome and deuterostome peptides.

Second, all sequences display two cysteine residues that can form a disulfide bond. Notably, the size of the disulfide loop appears conserved in all protostomes (4 residues including the WK motif) but varies in deuterostomes: while UII and URP share the loop size with protostome SSRP-like peptides, SS and CST display extended loops with 10 or 12 residues in chordates and starfish, respectively (**Figure 4**). If, as we propose here, the protostome and deuterostome systems are indeed related, following multiple rounds of whole genome duplication in chordates, UII and URP retained the ancestral loop size while SS and CST did not. We suggest that loop size correlates with structural stability but does not directly correlate with biological activity. Shorter loop sizes have been successfully introduced in therapeutic SS analogs to improve the *in vivo* half-life of these analogs while retaining and in some cases improving biological activity at some SS receptor subtypes (Pless 2005).

Third, when screened against the five subtypes of the human SS receptors, the neuroendocrine molluscan SSRP-like peptide exhibited micromolar activity at the human SST_2_, indicating that the ligands of the human and molluscan receptors are functionally related (**Figure S5**).

Fourth, mining the genome of the model annelid *C. teleta* for an SS-like receptor led to the discovery of a receptor that clusters with human SSRP receptors and that, when expressed in human HEK cells and tested using the PRESTO-Tango assay, is potently activated by the *C. teleta* SSRP-like peptide 1 identified here (EC_50_ = 13.1 nM).

Finally, the ability of consomatins from some fish-hunting cone snails to activate various subtypes of the human SS receptor (Ramiro et al. 2021) strongly suggests that the molluscan, and thereby protostome SSRPs-like peptides are not only related to chordate SSRPs but can be adapted to potently and selectively activate the chordate SS receptors.

Even though these lines of evidence strongly suggest a common ancestry of chordate SSRPs and protostome SSRP-like peptides, we note that other observations are less conclusive. The gene structures of protostome SSRPs differ from those observed for members of chordate SSRPs. Genes of human SS and CST only have a single phase 0 intron, while those encoding UII and URP contain a single phase 0 intron and 2-3 additional phase 1 introns (**Figure S2**). The echinoderm SS-like genes mirror SS and CST with a single phase 0 intron (**Figure S2**). It was previously proposed that an N-terminal phase 0 intron is a conserved characteristic of the SSRP gene family (Zhang et al. 2020). While all protostome SSRP-like genes identified here indeed share the phase 0 intron with SS/CST and one phase 1 intron with UII/URP, considering the varying gene structure of the chordate hormones, this is only a weak indicator of common ancestry.

Additionally, while we were able to show potent activation of an annelid GPCR related to the chordate SS receptors upon application of an annelid SSRP-like peptide (**Figure 5**), we were not able to identify the endogenous target receptor of molluscan SSRP-like peptides. While we have tested several additional molluscan receptor candidates with sequence similarity to the chordate SS receptor and the annelid SSRP-like receptor, we could not detect any activation of these receptors by the molluscan or annelid SSRP-like peptide, or peptides of the human SSRP family (data not shown). Whether the lack of activity is due to the type of assay used here, the identity and biology of the proposed target receptors, or to an incorrectly predicted mature SSRP-like peptide remains to be determined.

However, if, as we propose here, the protostome SSRP-like signaling peptides and consomatins are indeed related to chordate SSRPs, some of these peptides could potentially be new ligands of chordate receptors known to be activated by members of the SSRP family, namely the SST_1-5_, the UT, the ghrelin receptor and the MRGPRX2. This may particularly be the case for members of the highly diversified consomatin family. Previously, we showed that some consomatins from fish-hunters activate different subtypes of the human SS receptor and provide new drug leads for pain and neuroendocrine tumors (Ramiro et al. 2021). Here, we provide >500 additional consomatin sequences from diverse species of cone snails for future receptor target identification and biomedical exploration.

We note that the evolution of consomatins closely resembles that of cone snail venom insulins, toxins that induce dangerously low blood sugar by activating the insulin receptor of prey: once recruited into venom, cone snail insulins experience strong positive selection resulting in a hyper-diverse family of weaponized hormones (Safavi-Hemami et al. 2016). As observed for venom insulins, consomatins of fish-hunters tend to resemble the signaling SSRP-like sequences of fish. Likewise, consomatins expressed in worm hunters tend to be similar to those found in annelids. This pattern is consistent with the hypothesis that many if not all venom insulins and consomatins are used in prey capture, and as a consequence, they are strongly selected to interact with receptors that appear regularly in a snail’s diet. Furthermore, because of their streamlined role in predation, these weaponized hormones display certain properties that their signaling counterparts may not: unlike human insulin, weaponized insulins from *Conus tulipa* and *C. geographus* act rapidly because they do not self-associate into dimers and hexamers (Ahorukomeye et al. 2019; Menting et al. 2016), and, unlike human somatostatin, consomatins from *C. rolani* and *C. geographus* are highly stable and selective for certain SST receptor subtypes (Ramiro et al. 2021). Thus, weaponized hormones evolved advantageous properties that can inform on the design of new drug leads for diseases associated with native human hormones (Ramiro et al. 2021; Xiong et al. 2021; Xiong et al. 2020).

Additionally, we provide a set of new SSRP-like signaling sequences from various bilaterian phyla, including some invertebrate animal models such as *P.dumerilli*. Based on several observations we believe that this signaling system not only exists in these animals but plays important physiological roles. First, SSRP-like sequences were recovered from several transcriptome datasets demonstrating that these genes are actively expressed. Second, all these sequences have an N-terminal signal sequence and are therefore secreted to likely function in neuroendocrine signaling events. One member of the SSRP-like family, namely apUII from *A. californica*, has already been shown to be secreted in the nervous system where it exerts inhibitory effects on motor programs involved in feeding (Romanova et al. 2012). Others are likely to also serve important functions. Third, the broad presence and targeted use of consomatins by worm-hunting cone snails strongly suggests an important functional role of the SSRP-like signaling systems in annelids. Lastly, the existence and expression of an annelid SS-like receptor (Ct-SSTL1) that can be potently activated by the *C. teleta* SSRP-like 1 peptide further supports an important physiological role of this signaling system. Understanding the functional role of these peptides in protostomes may provide new opportunities to study the SSRP-like signaling system in invertebrate model organisms, such as *C. teleta* and *A. californica*. Similarly, elucidating the function of SSRP-like peptides in xenacoelomorhps may provide fundamental insight into the biological role of the ancestral SSRP-like signaling system in animals which may ultimately expand our current understanding of the diverse physiological roles of SSRP in humans.

Finally, we provide a new paradigm for the discovery of novel signaling systems based on peptide toxins that specifically evolved to target such systems. With the growing availability of large peptide toxin databases by transcriptome and genome sequencing we anticipate that this approach will be used to identify other important signaling system in the future.

## Acknowledgements

We thank Kasper Kildegaard Sørensen for help with peptide synthesis, Noel Saguil for assistance with field collections, Maren Watkins for insightful discussions, and the High Throughput Genomics Core Facility at the University of Utah, USA for library preparation and transcriptome sequencing.

## Materials and Methods

### Transcriptome sequencing

Specimens of *C. textile*, *C. striatus* and *C. imperialis* were collected in Oahu, Hawai, USA. *C. rolani* and *C. geographus* specimens were collected in Cebu, Philippines under the Department of Agriculture – Bureau of Fisheries and Aquatic Resources-issued gratuitous permit no. GP-0084-15. Total RNA was extracted from the circumoesophageal nerve rings, salivary gland, venom bulbs, and foot using the Direct-zol RNA extraction kit (Zymo Research), with on-column DNase treatment, according to the manufacturer’s instructions. Library preparation and sequencing were performed by the University of Utah High Throughput Genomics Core Facility as previously described (Ahorukomeye et al. 2019). Briefly, paired-end sequencing was performed on an Illumina HiSeq2500 or NovaSeq 6000 instrument. Adapter trimming of de-multiplexed raw reads was performed using fqtrim (v0.9.4), followed by quality trimming and filtering using prinseq-lite (Schmieder and Edwards 2011). Error correction was performed using the BBnorm ecc tool, part of the BBtools package (open source software, Joint Genome Institute). Trimmed and error-corrected reads were assembled using Trinity (version 2.2.1) (Grabherr et al. 2011; Haas et al. 2013) with a k-mer length of 31 and a minimum k-mer coverage of 10. Expression levels were calculated as transcripts per million (tpm). Tpm values were calculated using the RSEM program (Li and Dewey 2011). (**Supporting File 1**).

### Transcriptome assemblies and analyses

Raw sequencing data of publicly available cone snail venom gland transcriptomes were retrieved from the NCBI, DDBJ and the CNGB sequence repositories, and assembled using the same methods as described for tissues sequenced in this study. The list of all transcriptomes used in this study is provided in **Supporting File 1**. Consomatin sequences were identified from these datasets based on tblastn with sequences from (Ramiro et al. 2021) using e-value of 0.1. Sequences with the C-terminal C.WK.C-motif were identified as consomatins.

### Gene structure analysis

The location, size, and phases of introns were identified using the online version of Splign (Kapustin et al. 2008). Signal sequences were calculated using either command-line or online versions of SignalP 5.0 (Almagro Armenteros et al. 2019).

### Phylogenetic analysis

Multiple sequence alignments of protein sequences were performed using MAFFT v 7.310 (Katoh and Standley 2013). The tree in **Figure 3A** was constructed using a Bayesian analysis of phylogeny with Mr. Bayes v.3.2.6 (Huelsenbeck and Ronquist 2001; Ronquist and Huelsenbeck 2003) with mixed amino acid models. The analysis was performed with two runs of four Markov chains for 1,000,000 generations at which point the two chains had converged. The first 25 % of samples were discarded as burn-in. The tree uses the signaling SSRP-like precursor from *Aplysia californica* as an outgroup. The tree in **Figure 5A** and **S3** was constructed using maximum likelihood with the program IQ-TREE v1.6.1 (Chernomor et al. 2016; Nguyen et al. 2015). The program chose the model of evolution according to the Bayesian information criterion, which was LG+F+I+G4. The tree was rooted with human galanin and kisspeptin receptors, which form a known outgroup of the SSRP receptors.

### Selection analysis

Signaling SSRP-like genes were extracted from the nerve ring transcriptomes of *C. rolani, C. geographus, C. textile,* and *C. imperialis* and from the genomes of *C. ventricosus,* and *C. betulinus.* Twenty random toxin genes were obtained from NCBI GenBank (Supporting file 2). The three groups of the toxin genes and the two signaling genes were all analyzed separate and identically as follows: First, the amino acid sequences were aligned using ClustalO v 1.2.4 (Sievers et al. 2011). The amino acid alignment was then converted to the corresponding nucleotide alignment using the program PAL2NAL (Suyama et al. 2006) with the flag ‘-nogap’. The synonymous and non-synonymous substitutions were calculated from this codon alignment with the program codeml (v4.9) from PAML (Yang 2007) under the F3×4 model of evolution. We calculated the likelihood of several different models of evolution, including pairwise comparisons of all sequences, a beta distribution of dN/dS between 0 and 1 (M7), and a beta distribution of dN/dS with positive selection at some sites (M8). Conventional likelihood ratio tests were used by comparing the M7 and M8. The sites with positive selection were estimated with the Bayes empirical Bayes analysis.

The conservation scores for the different groups (signaling SSRP-like and consomatins) were calculated with the program rate4site (Mayrose et al. 2004) using an alignment generated by ClustalO v 1.2.4 and a maximum likelihood tree generated by IQ-tree using the same alignment. The conservation score for each site is visualized with a sliding window of 5 amino acids in the figure.

### Consomatin exon recovery and principal component analysis

The SRA datasets of cone snails from Bioproject PRNJ526781 were downloaded from NCBI in September 2021 with fastq-dump 2.8.2 (SRA toolkit development team https://trace.ncbi.nlm.nih.gov/Traces/sra/sra.cgi?view=software). The individual dataset was then preprocessed with fastp v.0.20.1 (Chen et al. 2018) to filter out low quality reads and were converted to fasta-format with EMBOSS seqret v. 6.6.0.0 (Madeira et al. 2019). To identify reads encoding consomatins we used the toxin sequences from earlier identified consomatins as queries in a tblastn search against the database using an e-value of 10. The hits were then assembled using Trinity v.2.13.2 (Grabherr et al. 2011; Haas et al. 2013) as single reads. The generated contigs were translated in all reading frames using EMBOSS getorf v.6.6.0.0 (Madeira et al. 2019). The putative toxins were extracted with the regular expression ‘[^C]{0,2}C[^C]WK[^C]C[^RKC]{0,6}’ and only contigs with 5-fold coverage were kept for the subsequent analyses. The pool of extracted toxins from the SRA-data were used as tblastn-queries in two additional iterations of toxin identification.

In addition to the identified toxins, we also used signaling SSRP-like peptides from annelids, molluscs and vertebrates in the analysis. These were modified to only have two amino acid residues preceding the cysteine-ring to match the structure of the toxin-sequences in this analysis.

We extracted the length, molecular weight, aromaticity, and isoelectric point, as well as the count for the 150 most common 1-3mers across all the sequences. Principal components were calculated using the sklearn library v 0.24.1 (Pedregosa et al. 2011) in Python 3.6.9 on data scaled to remove the mean and unit variance. The two first principal components were used to visualize the sequences using R 3.6.3 with the library tidyverse 1.3.1 (Wickham et al. 2019).

### Peptide synthesis

Ct-SSRPL1 predicted from the whole-body transcriptome of the annelid *C. teleta* was synthesized by solid-phase peptide synthesis (SPPS) in 0.1 mmol scale using preloaded Fmoc-Trp Tentagel R HMPA resin from Rapp Polymere (Tuebingen, Germany). The sequence was H-ZASSGAQICIWKVCPPTPW-OH, where Z represents a pyroglutamic acid and Cys9 and Cys14 are connected by a disulfide bond. Fmoc-protected amino acids, coupling reagents and solvents used for the synthesis were purchased from Iris Biotech (Marktredwitz, Germany). The synthesis was performed on a Syro I instrument from Biotage (Uppsala, Sweden). Coupling conditions were room temperature for 2 × 120 min using 5.2 eq amino acids, 4.7 eq *N*-[(1*H*-benzotriazol-1-yl)(dimethylamino)methylene]-*N*-methylmethanaminium hexafluorophosphate *N*-oxide (HBTU), 5.2 eq 1-hydroxy-7-azabenzotriazol (HOAt), and 8 eq N,N-diisopropylethylamine (DIPEA) in dimethylformamide (DMF) relative to resin. Deprotection was performed at room temperature with 40 % piperidine in DMF for 3 min followed by 20 % piperidine in DMF for 15 min. Washing steps were with 2 × N-methyl-2-pyrrolidone (NMP), 1 × dichloromethane (DCM), and 1 × DMF. After completion of the peptide assembly, the peptidyl-resin was washed with 3 × DCM and dried. The peptide was released from approximately 0.05 mmol of resin using a mixture (3 mL) of 95% trifluoroacetic acid (TFA), 2.5 % triethylsilyl (TES), and 2.5 % water over 2.5 h. Cold diethylether (15 mL at −20 °C) was added and the mixture further cooled to −85 °C for 30 min before the peptide was isolated by centrifugation. The peptide was redissolved in a mixture of acetic acid, water, and acetonitrile (ACN) in volume ratios of 1:10:4 and freeze dried to remove unwanted carboxylation of Trp. The freeze dried compound was resuspended in 1 % vol/vol TFA and water. 0.1 M phosphate buffer and appr. ACN (3 mL) was added to 10 mL to this mixture. The pH was adjusted to 6 using 1 M hydrochloride acid (HCl). Dimethyl sulfoxide (DMSO, 1.0 mL) was added (10 % V/V) and the mixture was stirred for 20 h at room temperature to allow disulfide bond formation. The mixture was filtered and directly purified on a Dionex Ultimate 3000 HPLC system (Thermo Fisher, Waltham, USA) using a Luna C18(2) column from Phenomenex (Torrance, USA, 5 μm, 100 Å, 250 × 10 mm). A gradient of 5% - 100 % ACN in water/0.1 % formic acid was applied. The product was isolated and lyophilized by freeze drying to yield appr. 3.5 mg (0.0017 mmol). The final product was analyzed by liquid chromatography (LC)-mass spectrometry (MS) on a Dionex Ultimate 3000 ultra high performance LC (UHPLC) system from Thermo Fisher connected to an Impact HD mass spectrometer (Bruker, Bremen, Germany). The calculated monoisotopic MH^+1^ of the purified product was 2052.97 (detected monoisotopic MH^+1^: 2053.00) (**Figure S4A**).

Cr-SSRPL1 predicted from the nerve ring transcriptome of the cone snail *C. rolani* was custom synthesized by GenScript (Leiden, Netherlands). The sequence was H-SGGMSLCLWKVCPAAPWLVS-OH, where Cys7 and Cys12 are connected by a disulfide bond. The folded peptide was purified to > 95% by HPLC analysis and verified by MS. The calculated monoisotopic MH^+1^ of the purified product was 2103.02 (detected monoisotopic MH^+1^: 2103.39) (**Figure S4B**).

### GPCR assays

Unless otherwise stated, all chemicals used for cell culture were from Sigma-Aldrich (Burlington, MA, USA), and all medium and buffers are from ThermoFisher Scientific (Waltham, MA, USA).

HTLA cells (a kind gift from Prof. Hans Bräuner-Osborne) were maintained in DMEM supplemented with 10 % FBS (Biowest, Nuaillé, Franc), 100 U/mL penicillin and 100 μg/mL streptomycin, as well as 100 μg/mL hygromycin B and 2 μg/mL puromycin (growth medium) and detached by washing with Ca^2+^- and Mg^2+^-free PBS (Substrate Department, University of Copenhagen, DK). One million cells per well were seeded in 6 well plates in 2 mL growth medium the day before transfection and transfected with PolyFect (Qiagen, Düsseldorf, DE) using the manufacturers protocol (changing the medium to growth medium without hygromycin B and puromycin prior to transfection). 15,000 transfected cells per well were seeded in poly-D-lysine coated white clear bottom 384 well plates (Corning New York, NY, USA) in 40 μL DMEM supplemented with 1 % dFBS (ThermoFisher) (assay medium) and incubated overnight. The next day, medium was changed to 40 μL fresh assay medium, and compounds were added at 5 × concentration in 10 μL HBSS supplemented with 1 mM CaCl_2_, 1 mM MgCl_2_, 20 mM HEPES, pH adjusted to 7.4 with 10 M NaOH (assay buffer) with 0.1 % BSA added and sterile (0.2 μM) filtered, and cells were incubated overnight. The next day, the compounds were removed, and the cells incubated for 20 minutes in 20 μL of assay buffer with 0.01 % pluronic F-68 (ThermoFisher) and 5 % BrightGlo (Promega, Madison, WI, USA) before reading luminescence on a Molecular Devices SpectraMax iD5 with a 1 second integration time. Data was analyzed using GraphPad Prism 9 (San Diego, CA, USA). Concentration response curves were fitted to a four-parameter dose-response curve.

## Supporting Figures

**Figure S1.**
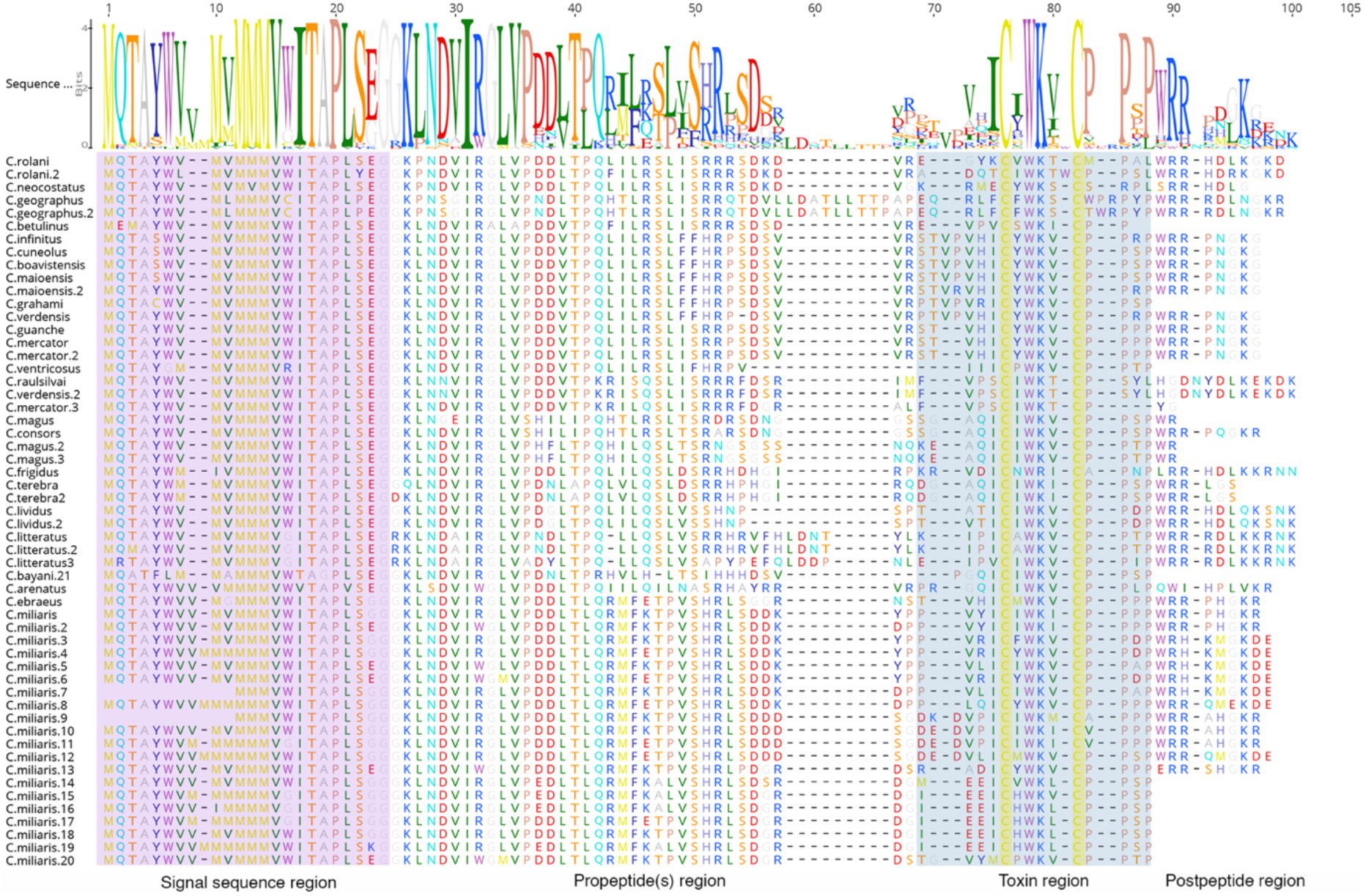
Multiple sequence alignment of consomatins identified in the venom gland transcriptomes of diverse cone snail species. Color coding as shown in Figure 2. Sequence logos were created using Geneious software (version R11).

**Figure S2.**
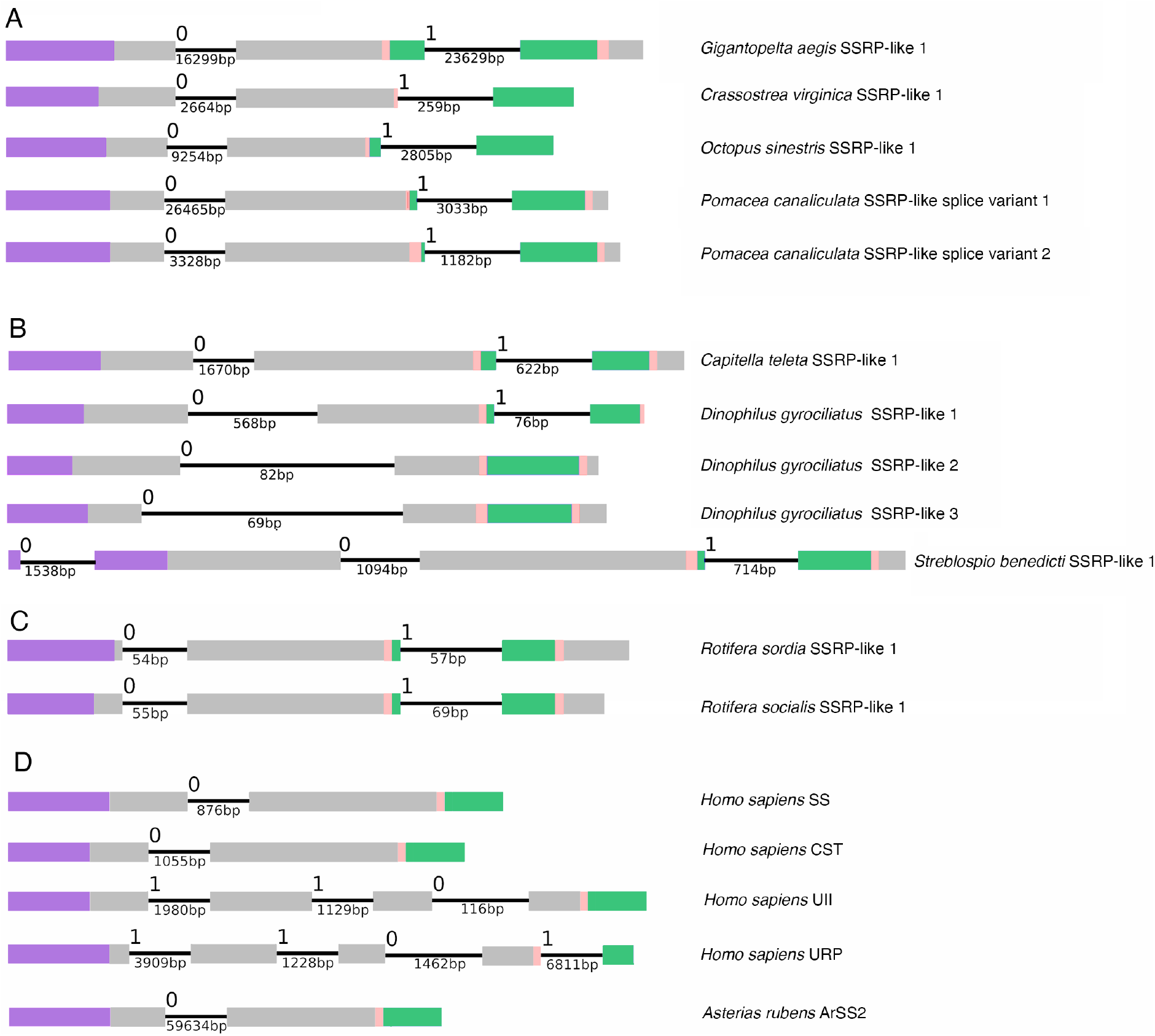
Gene structures of SSRP-like genes in (**A**) Mollusca, (**B**) Annelida, (**C**) Rotifera, and (**D**) Deuterostomes (human and starfish). Colors are according to figure 2; signal sequence (purple), propeptides (gray), mature toxin region (blue), mature signaling SSRP-like peptide region (green), and postpeptides (gray).

**Figure S3.**
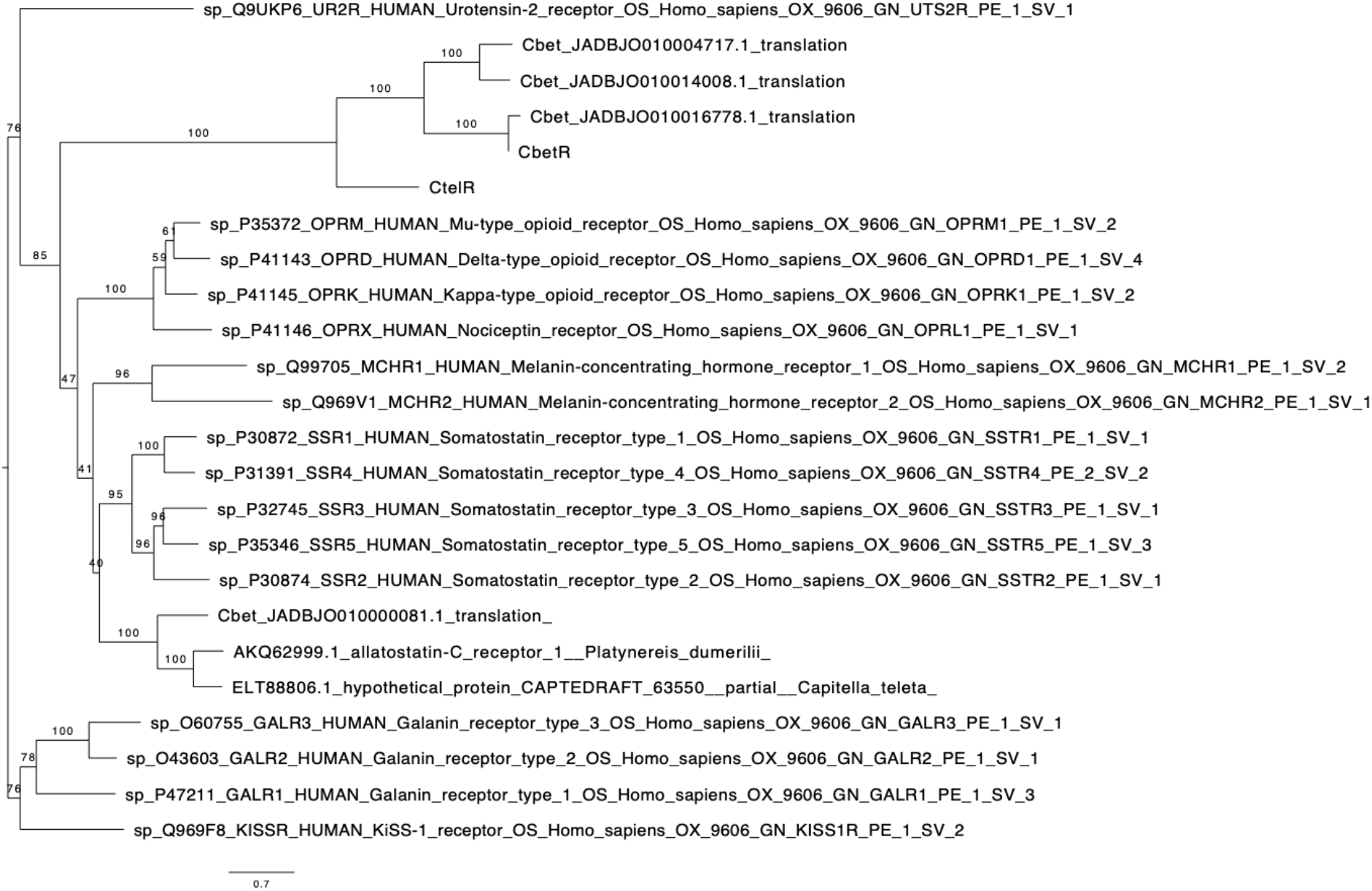
Maximum likelihood tree of SSRP-receptors from human, AST-C receptor from *Platynereis dumerilii* and putative receptors from *Conus betulinus* and *Capitella teleta*. The tree is a full version of **Figure 5A** in the main text. The bootstrap values were calculated with 1000 replicated from IQ-Tree’s UF-bootstrap method.

**Figure S4.**
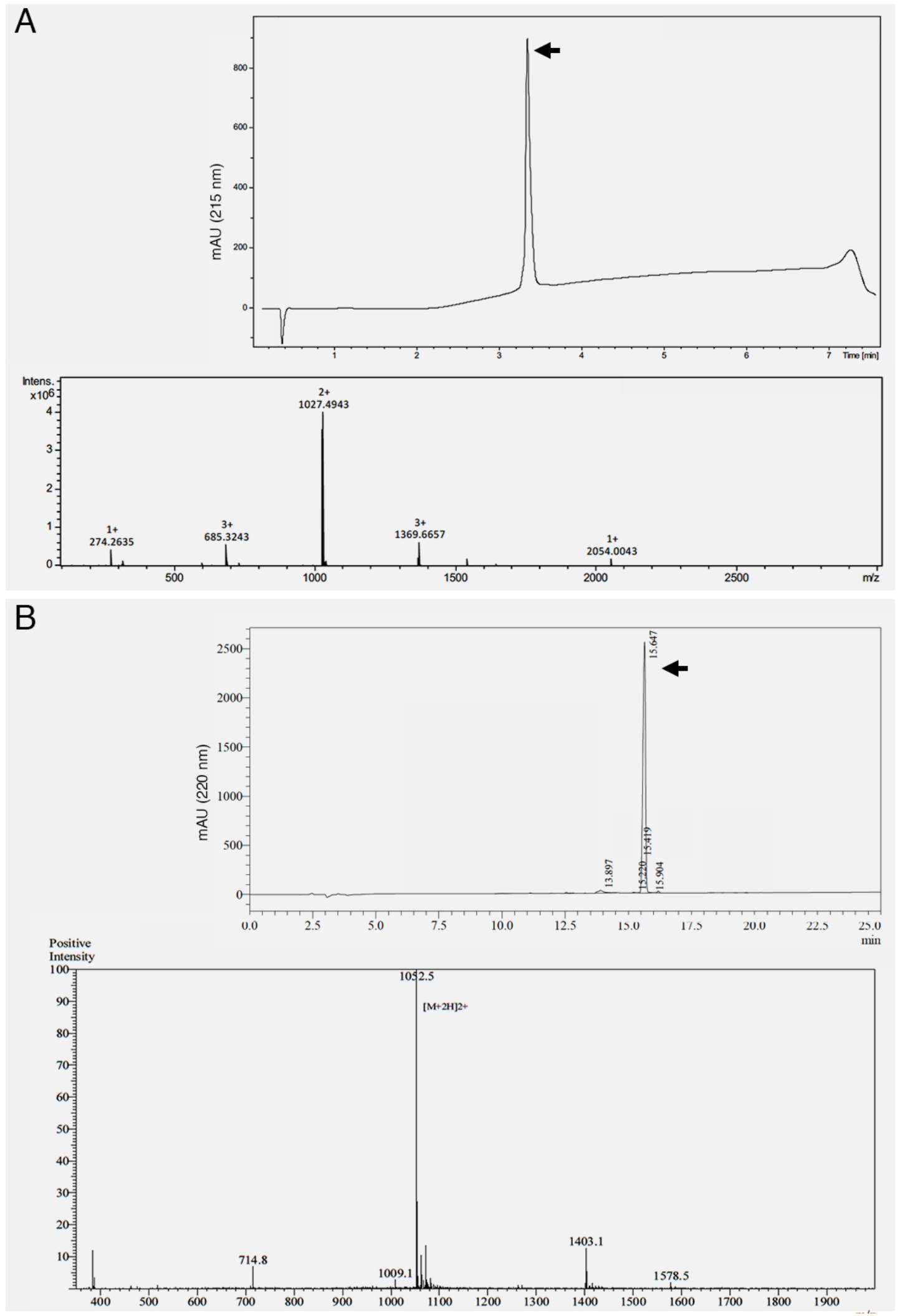
Quality control of synthetic (**A**) Ct-SSRPL1 from the annelid *Capitella teleta* and (**B**) Cr-SSRPL1 from *Conus rolani*. Peptides were synthesized using solid-phase synthesis as described in material and methods. The purity and integrity of the peptides was analyzed by reversed-phase high performance liquid chromatography (HPLC, upper panel) and mass spectrometry (MS/MS, lower panel) of the main HPLC peaks (black arrows).

**Figure S5.**
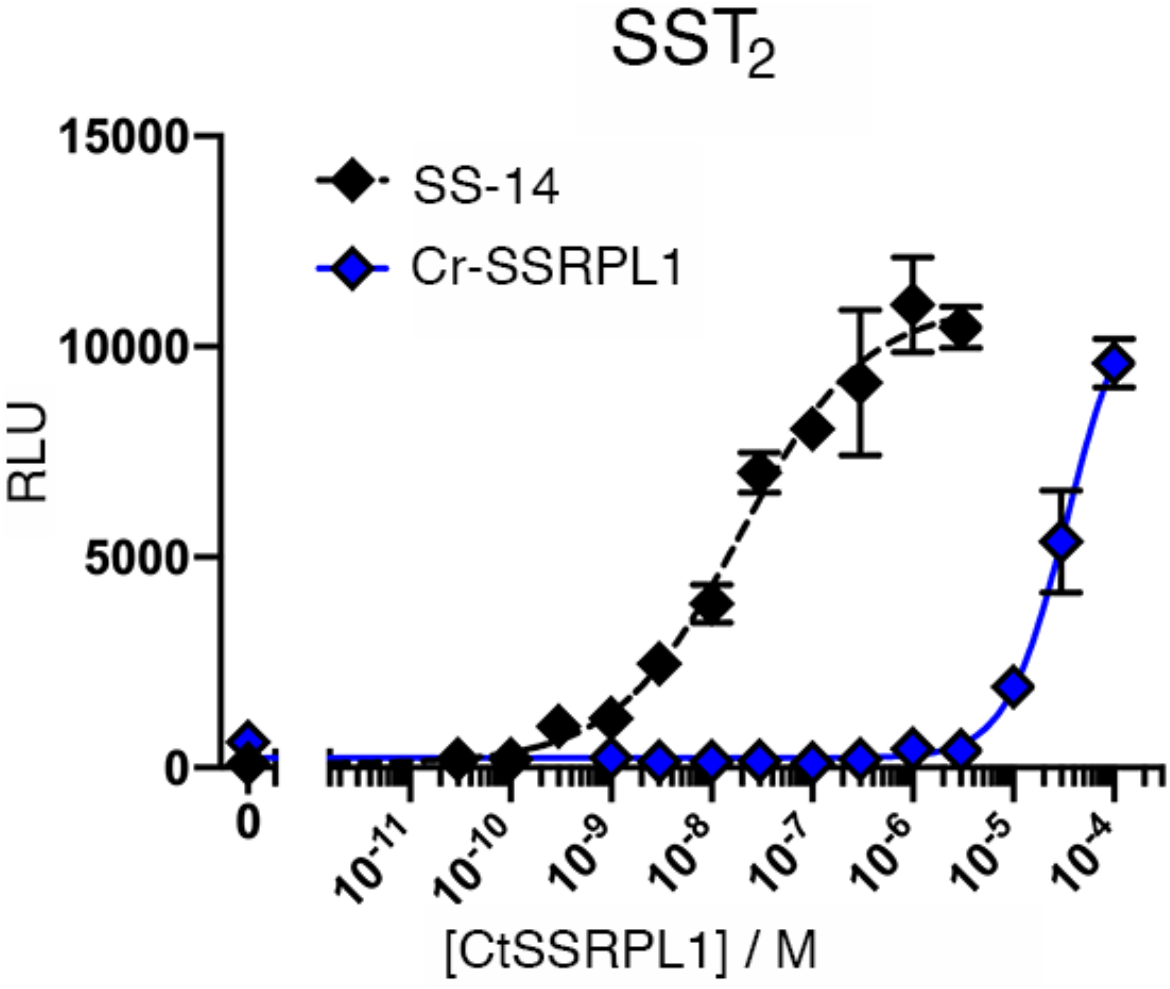
Concentration-response curve of human somatostatin-14 (SS) and the *Conus rolani* nerve ring SSRP-like peptide (Cr-SSRPL1) at the human somatostatin receptor 2 (SST_2_) in the PRESTO-Tango assay (error bars represent SD), showing an activation by Cr-SSRPL1 at micromolar concentrations.

## References

Ahorukomeye P, Disotuar MM, Gajewiak G, Karanth S, Watkins M, Robinson SD, Flórez Salcedo P, Smith NA, Smith BJ, Schlegel A et al. 2019. Fish-hunting cone snail venoms are a rich source of minimized ligands of the vertebrate insulin receptor. eLIFE. Feb 14, 2019.

Almagro Armenteros JJ, Tsirigos KD, Sønderby CK, Petersen TN, Winther O, Brunak S, von Heijne G, Nielsen H. 2019. Signalp 5.0 improves signal peptide predictions using deep neural networks. Nature Biotechnology. 37(4):420–423.

Aman JW, Imperial JS, Ueberheide B, Zhang MM, Aguilar M, Taylor D, Watkins M, Yoshikami D, Showers-Corneli P, Safavi-Hemami H et al. 2015. Insights into the origins of fish hunting in venomous cone snails from studies of conus tessulatus. Proc Natl Acad Sci U S A. 112(16):5087–5092.

Brazeau P, Vale W, Burgus R, Ling N, Butcher M, Rivier J, Guillemin R. 1973. Hypothalamic polypeptide that inhibits the secretion of immunoreactive pituitary growth hormone. Science. 179(4068):77–79.

Cannon JT, Vellutini BC, Smith J, Ronquist F, Jondelius U, Hejnol A. 2016. Xenacoelomorpha is the sister group to nephrozoa. Nature. 530(7588):89–93.

Chen S, Zhou Y, Chen Y, Gu J. 2018. Fastp: An ultra-fast all-in-one fastq preprocessor. Bioinformatics. 34(17):i884–i890.

Chernomor O, von Haeseler A, Minh BQ. 2016. Terrace aware data structure for phylogenomic inference from supermatrices. Systematic Biology. 65(6):997–1008.

Cruz LJ, de Santos V, Zafaralla GC, Ramilo CA, Zeikus R, Gray WR, Olivera BM. 1987. Invertebrate vasopressin/oxytocin homologs. Characterization of peptides from *conus geographus* and *conus striatus* venoms. The Journal of biological chemistry. 262(33):15821–15824.

Deghenghi R, Papotti M, Ghigo E, Muccioli G. 2001. Cortistatin, but not somatostatin, binds to growth hormone secretagogue (ghs) receptors of human pituitary gland. J Endocrinol Invest. 24(1):Rc1–3.

Duda TF, Jr., Kohn AJ, Palumbi S. 2001. Origins of diverse feeding ecologies within *conus*, a genus of venomous marine gastropods. Biological Journal of the Linnean Society. 73(4):391–409.

Elphick MR, Mirabeau O, Larhammar D. 2018. Evolution of neuropeptide signalling systems. The Journal of experimental biology. 221(Pt 3).

Grabherr MG, Haas BJ, Yassour M, Levin JZ, Thompson DA, Amit I, Adiconis X, Fan L, Raychowdhury R, Zeng Q et al. 2011. Full-length transcriptome assembly from rna-seq data without a reference genome. Nat Biotechnol. 29(7):644–652.

Grimmelikhuijzen CJP, Hauser F. 2012. Mini-review: The evolution of neuropeptide signaling. Regul Pept. 177 Suppl:S6–9.

Haas BJ, Papanicolaou A, Yassour M, Grabherr M, Blood PD, Bowden J, Couger MB, Eccles D, Li B, Lieber M et al. 2013. De novo transcript sequence reconstruction from rna-seq using the trinity platform for reference generation and analysis. Nature Protocols. 8(8):1494–1512.

Hejnol A, Pang K. 2016. Xenacoelomorpha’s significance for understanding bilaterian evolution. Current Opinion in Genetics & Development. 39:48–54.

Huelsenbeck JP, Ronquist F. 2001. Mrbayes: Bayesian inference of phylogenetic trees. Bioinformatics. 17(8):754–755.

Jékely G. 2013. Global view of the evolution and diversity of metazoan neuropeptide signaling. Proceedings of the National Academy of Sciences. 110(21):8702–8707.

Jondelius U, Raikova OI, Martinez P. 2019. Xenacoelomorpha, a key group to understand bilaterian evolution: Morphological and molecular perspectives. In: Pontarotti P, editor. Evolution, origin of life, concepts and methods. Cham: Springer International Publishing. p. 287–315.

Kapli P, Telford MJ. 2020. Topology-dependent asymmetry in systematic errors affects phylogenetic placement of ctenophora and xenacoelomorpha. Science Advances. 6(50):eabc5162.

Kapustin Y, Souvorov A, Tatusova T, Lipman D. 2008. Splign: Algorithms for computing spliced alignments with identification of paralogs. Biology Direct. 3(1):20.

Katoh K, Standley DM. 2013. Mafft multiple sequence alignment software version 7: Improvements in performance and usability. Molecular biology and evolution. 30(4):772–780.

Kohn AJ. 1956. Piscivorous gastropods of the genus *conus*. Proceedings of the National Academy of Sciences of the United States of America. 42(3):168–171.

Li B, Dewey CN. 2011. Rsem: Accurate transcript quantification from rna-seq data with or without a reference genome. BMC Bioinformatics. 12(1):323.

Li Q, Barghi N, Lu A, Fedosov AE, Bandyopadhyay PK, Lluisma AO, Concepcion GP, Yandell M, Olivera BM, Safavi-Hemami H. 2017. Divergence of the venom exogene repertoire in two sister species of turriconus. Genome biology and evolution. 9(9):2211–2225.

Madeira F, Park YM, Lee J, Buso N, Gur T, Madhusoodanan N, Basutkar P, Tivey ARN, Potter SC, Finn RD et al. 2019. The embl-ebi search and sequence analysis tools apis in 2019. Nucleic Acids Res. 47(W1):W636–W641.

Mair GR, Halton DW, Shaw C, Maule AG. 2000. The neuropeptide f (npf) encoding gene from the cestode, moniezia expansa. Parasitology. 120(1):71–77.

Martín-Durán JM, Vellutini BC, Marlétaz F, Cetrangolo V, Cvetesic N, Thiel D, Henriet S, Grau-Bové X, Carrillo-Baltodano AM, Gu W et al. 2021. Conservative route to genome compaction in a miniature annelid. Nat Ecol Evol. 5(2):231–242.

Martinez V. 2015. Handbook of biologically active peptides. copyright © 2013 elsevier inc. All rights reserved. In: Takei Y, Ando H, Tsutsui K, editors. Elsevier. p. 1320–1329.

Mayrose I, Graur D, Ben-Tal N, Pupko T. 2004. Comparison of site-specific rate-inference methods for protein sequences: Empirical bayesian methods are superior. Molecular biology and evolution. 21(9):1781–1791.

Menting JG, Gajewiak J, MacRaild CA, Chou DH, Disotuar MM, Smith NA, Miller C, Erchegyi J, Rivier JE, Olivera BM et al. 2016. A minimized human insulin-receptor-binding motif revealed in a conus geographus venom insulin. Nature structural & molecular biology. 23(10):916–920.

Mirabeau O, Joly J-S. 2013. Molecular evolution of peptidergic signaling systems in bilaterians. Proceedings of the National Academy of Sciences. 110(22):E2028–E2037.

Moller LN, Stidsen CE, Hartmann B, Holst JJ. 2003. Somatostatin receptors. Biochimica et biophysica acta. 1616(1):1–84.

Nguyen L-T, Schmidt HA, von Haeseler A, Minh BQ. 2015. Iq-tree: A fast and effective stochastic algorithm for estimating maximum-likelihood phylogenies. Molecular biology and evolution. 32(1):268–274.

Nybakken J, Perron FE. 1988. Ontogenetic change in the radula of *conus magus* (gastropoda). Marine Biology. 98:239–242.

Olivera BM, Seger J, Horvath MP, Fedosov AE. 2015. Prey-capture strategies of fish-hunting cone snails: Behavior, neurobiology and evolution. Brain, behavior and evolution. 86(1):58–74.

Ong KL, Wong LY, Cheung BM. 2008. The role of urotensin ii in the metabolic syndrome. Peptides. 29(5):859–867.

Pedregosa F, Varoquaux Ge l, Gramfort A, Michel V, Thirion B, Grisel O, Blondel M, Prettenhofer P, Weiss R, Dubourg V et al. 2011. Scikit-learn: Machine learning in python. Journal of Machine Learning Research. 12(Oct):2825––2830.

Phuong MA, Alfaro ME, Mahardika GN, Marwoto RM, Prabowo RE, von Rintelen T, Vogt PWH, Hendricks JR, Puillandre N. 2019. Lack of signal for the impact of conotoxin gene diversity on speciation rates in cone snails. Syst Biol. 68(5):781–796.

Pless J. 2005. The history of somatostatin analogs. J Endocrinol Invest. 28(11 Suppl International):1–4.

Puillandre N, Bouchet P, Duda TF, Jr., Kauferstein S, Kohn AJ, Olivera BM, Watkins M, Meyer C. 2014. Molecular phylogeny and evolution of the cone snails (gastropoda, conoidea). Molecular phylogenetics and evolution. 78:290–303.

Ramiro IBL, Bjørn-Yoshimoto WE, Imperial JS, Gajewiak J, Watkins M, Taylor D, Resager W, Ueberheide B, Bräuner-Osborne H, Whitby FG et al. 2021. Somatostatin venom analogs evolved by fish-hunting cone snails: From prey capture behavior to identifying drug leads. bioRxiv.2021.2010.2026.465842.

Robas N, Mead E, Fidock M. 2003. Mrgx2 is a high potency cortistatin receptor expressed in dorsal root ganglion. J Biol Chem. 278(45):44400–44404.

Romanova EV, Sasaki K, Alexeeva V, Vilim FS, Jing J, Richmond TA, Weiss KR, Sweedler JV. 2012. Urotensin ii in invertebrates: From structure to function in aplysia californica. PloS one. 7(11):e48764.

Ronquist F, Huelsenbeck JP. 2003. Mrbayes 3: Bayesian phylogenetic inference under mixed models. Bioinformatics. 19(12):1572–1574.

Safavi-Hemami H, Lu A, Li Q, Fedosov AE, Biggs J, Showers Corneli P, Seger J, Yandell M, Olivera BM. 2016. Venom insulins of cone snails diversify rapidly and track prey taxa. Molecular biology and evolution. 33(11):2924–2934.

Schmieder R, Edwards R. 2011. Quality control and preprocessing of metagenomic datasets. Bioinformatics. 27(6):863–864.

Semmens DC, Mirabeau O, Moghul I, Pancholi MR, Wurm Y, Elphick MR. 2016. Transcriptomic identification of starfish neuropeptide precursors yields new insights into neuropeptide evolution. Open biology. 6(2):150224.

Sievers F, Wilm A, Dineen D, Gibson TJ, Karplus K, Li W, Lopez R, McWilliam H, Remmert M, Söding J et al. 2011. Fast, scalable generation of high-quality protein multiple sequence alignments using clustal omega. Molecular Systems Biology. 7(1):539.

Suyama M, Torrents D, Bork P. 2006. Pal2nal: Robust conversion of protein sequence alignments into the corresponding codon alignments. Nucleic Acids Res. 34(Web Server issue):W609–612.

Tostivint H, Ocampo Daza D, Bergqvist CA, Quan FB, Bougerol M, Lihrmann I, Larhammar, D. 2014. Molecular evolution of gpcrs: Somatostatin/urotensin ii receptors. Journal of molecular endocrinology. 52(3):T61–86.

Vaudry H, Do Rego JC, Le Mevel JC, Chatenet D, Tostivint H, Fournier A, Tonon MC, Pelletier G, Conlon JM, Leprince J. 2010. Urotensin ii, from fish to human. Ann N Y Acad Sci. 1200:53–66.

Veenstra JA. 2009. Allatostatin c and its paralog allatostatin double c: The arthropod somatostatins. Insect Biochem Mol Biol. 39(3):161–170.

Wickham H, Averick M, Bryan J, Chang W, McGowan L, François R, Grolemund G, Hayes A, Henry L, Hester J et al. 2019. Welcome to the tidyverse. Journal of Open Source Software. 4:1686.

Woodward SR, Cruz LJ, Olivera BM, Hillyard DR. 1990. Constant and hypervariable regions in conotoxin propeptides. Embo Journal. 9(4):1015–1020.

Xiong X, Blakely B, Kim JH, Menting J, Schubert HL, R. A, T. G, I.B. S, Delaine C, Y.W. Z et al. 2021. Visualization of insulin receptor activation by a novel insulin analog with elongated a chain and truncated b chain. Nature Chemical Biology. in review.

Xiong X, Menting JG, Disotuar MM, Smith NA, Delaine CA, Ghabash G, Agrawal R, Wang X, He X, Fisher SJ et al. 2020. A structurally minimized yet fully active insulin based on cone-snail venom insulin principles. Nature structural & molecular biology. 27(7):615–624.

Yañez-Guerra LA, Zhong X, Moghul I, Butts T, Zampronio CG, Jones AM, Mirabeau O, Elphick MR. 2020. Echinoderms provide missing link in the evolution of prrp/snpf-type neuropeptide signalling. eLife. 9:e57640.

Yang Z. 2007. Paml 4: Phylogenetic analysis by maximum likelihood. Molecular biology and evolution. 24(8):1586–1591.

Zakas C, Harry ND, Scholl EH, Rockman MV. 2021. The genome of the poecilogonous annelid streblospio benedicti. bioRXiv.

Zhang M, Wang Y, Li Y, Li W, Li R, Xie X, Wang S, Hu X, Zhang L, Bao Z. 2018. Identification and characterization of neuropeptides by transcriptome and proteome analyses in a bivalve mollusc patinopecten yessoensis. Front Genet. 9:197–197.

Zhang Y, Yañez Guerra LA, Egertová M, Zampronio CG, Jones AM, Elphick MR. 2020. Molecular and functional characterization of somatostatin-type signalling in a deuterostome invertebrate. Open biology. 10(9):200172.

